# Transcriptome sequencing and bioinformatics analysis of kidney tissue in a tree shrew model of calcium oxalate nephrolithiasis

**DOI:** 10.1101/2024.02.18.580876

**Authors:** Guang Wang, Ziye Huang, Yuyun Wu, Rui Xu, Jiongming Li

## Abstract

**Background:** Kidney stones, predominantly composed of calcium oxalate, are a prevalent and recurrent urological condition. Given their high incidence and recurrence rates, understanding their pathogenesis and identifying effective treatment strategies are imperative.

**Methods:** In this study, we established a calcium oxalate nephrolithiasis model using tree shrews, a primate-like animal species. When compared to commonly used rodent models (rats and mice), the tree shrew model demonstrated superior reproducibility and relevance. And leveraging transcriptome sequencing and comprehensive bioinformatics analysis.

**Results:** we identified 1,927 differentially expressed genes, including 1,450 upregulated and 476 downregulated genes. Furthermore, we annotated these genes to 41 KEGG enriched pathways and 1,413 GO enrichments, encompassing 1276 Biological Processes, 72 Cellular Components, and Molecular Functions. Notably, we prioritized the top 50 core genes that could potentially underlie the pathogenesis of calcium oxalate nephrolithiasis.

**Conclusions:** Our findings establish the tree shrew as a relevant model for studying kidney stone formation and provide valuable insights into the underlying molecular mechanisms. These insights hold promise for the development of novel therapeutic strategies to address this significant health burden.

## Introduction

Kidney stones, a common urological condition, are characterized by their prevalence and propensity for recurrence. Reports suggest that approximately 10% of individuals are affected by kidney stones [1], with a recurrence rate nearing 10% within one year and exceeding 75% within 20 years [2]. Notably, calcium oxalate, a difficult-to-treat component by medicine, accounts for over 80% of kidney stones [3]. Current therapeutic approaches primarily rely on surgical intervention, despite limited progress in the field. The majority of existing research has focused on rodent models, such as rats and mice, raising questions about the translatability of these findings to primates and the subsequent development of effective treatment strategies.

The tree shrew, a primate-like mammal, offers a unique opportunity to address these challenges. Due to its remarkable physiological, biochemical, and genetic similarity to humans, the tree shrew has been successfully employed in diverse fields, including neuroscience, oncology, virology, and drug discovery. Particularly noteworthy is its utilization in virus infection and immune response studies, where it has demonstrated exceptional utility in elucidating virus pathogenesis and developing innovative treatment approaches. [4-7]

In our previous work, we established a calcium oxalate nephrolithiasis model using tree shrews and compared it to rodent models. The results were promising, indicating that the tree shrew model outperformed the rodent models in terms of relevance and reproducibility [8]. Leveraging this tree shrew model, we obtained kidney samples from both kidney stone-affected and control animals and performed transcriptome sequencing and comprehensive bioinformatics analysis. This approach allowed us to gain deeper insights into the molecular mechanisms underlying calcium oxalate nephrolithiasis, laying the foundation for the development of novel therapeutic strategies.

## Methods

### Data Preparation

Kidney tissue samples from tree shrews were shipped in dry ice to the sequencing facility at Qinglian Bai’ao Biotechnology Co., Ltd., Beijing, China. These samples underwent transcriptome sequencing on the Novaseq 6000 PE150 platform.

### Quality Control and Transcriptome Library Assessment

Using FastQC 0.10.1, the raw sequencing data was filtered to obtain high-quality reads (Clean Data). Subsequently, the Clean Data was mapped to the specific reference genome of Tupaia chinensis (Genome assembly TupChi_1.0, available at https://www.ncbi.nlm.nih.gov/datasets/genome/GCF_000334495.1/) using hisat2 2.1.0 and SAMtools 1.3.1. This alignment process evaluated the sequencing saturation, gene coverage, and overall alignment efficiency.

### Quantification of Gene Expression

The stringtie tool was employed to count the number of reads per gene. These read counts were then normalized using the Transcripts Per Million (TPM) method to mitigate biases arising from sequencing depth and sample composition.

### Differential Expression Analysis

Differential gene expression under various conditions was analyzed using statistical models implemented in R packages, notably limma. Genes exhibiting significant expression changes (p < 0.05) and an absolute fold change greater than 1 were designated as differentially expressed.

### Functional Annotation and Pathway Enrichment

Functional annotations, including Gene Ontology (GO) terms, were assigned to the differentially expressed genes using R packages like clusterProfiler and org.Hs.eg.db. Additionally, pathway enrichment analysis was conducted using the online tool STRING (https://string-db.org) to identify significantly enriched biological processes and signaling pathways.

### Protein-Protein Interaction (PPI) Network Analysis

The PPI network of the differentially expressed genes was constructed using the STRING database. This network was then imported into Cytoscape 3.8.2 with the cytoHubba plugin, employing the MCC algorithm to identify the top 50 most influential genes (hubs) within the network.

### Data Visualization and Reporting

Data visualization was achieved using the ggplot2 package in R, along with other specialized bioinformatics visualization tools like ComplexHeatmap. These visualizations provided intuitive representations of the sequencing data, differential expression patterns, and functional annotations.

## Results

### Differentially Expressed Genes

A total of 1927 genes were differentially expressed, with 1450 genes upregulated and 476 genes downregulated. The top 20 upregulated genes were CHIT1, GPNMB, DCSTAMP, MMP1, RIPPLY3, MMP12, CLEC4E, GLDN, SERPINA3, IL1RL1, GCG, CCL22, CUNH7orf25, SLAMF7, ITGAX, RBP4, SLC37A2, RGS1, TNFRSF18, and AP3B2. The top 20 downregulated genes were ADGRG7, CRIP3, ABCG8, OSR1, TAL2, FAM162B, GJD2, ARHGAP36, ERAS, LVRN, ERMN, DDIT4L, LRRN3, SHOX2, LMX1A, HDC, TUBAL3, TENM1, EMILIN3, and TAFA1. (Supplementary Table 1)

### Functional Annotation

The top 10 Biological Processes identified by GO enrichment analysis were Myeloid leukocyte activation, T cell differentiation, Cell activation involved in immune response, Regulation of leukocyte differentiation, Positive regulation of leukocyte activation, Lymphocyte differentiation, Regulation of T cell activation, Positive regulation of cell activation, Mononuclear cell differentiation, and Leukocyte cell−cell adhesion. The top 10 Cellular Components were Tertiary granule, External side of plasma membrane, Phagocytic vesicle, Specific granule, Collagen −containing extracellular matrix, Membrane raft, Membrane microdomain, Cytoplasmic vesicle lumen, Vesicle lumen, and Endocytic vesicle. The top 10 Molecular Functions were Chemokine binding, Cytokine receptor activity, Pattern recognition receptor activity, Immune receptor activity, Cytokine binding, Integrin binding, Glycosaminoglycan binding, Extracellular matrix structural constituent, Heparin binding, and Cytokine activity (Figure 1).

**Fig. 1.**
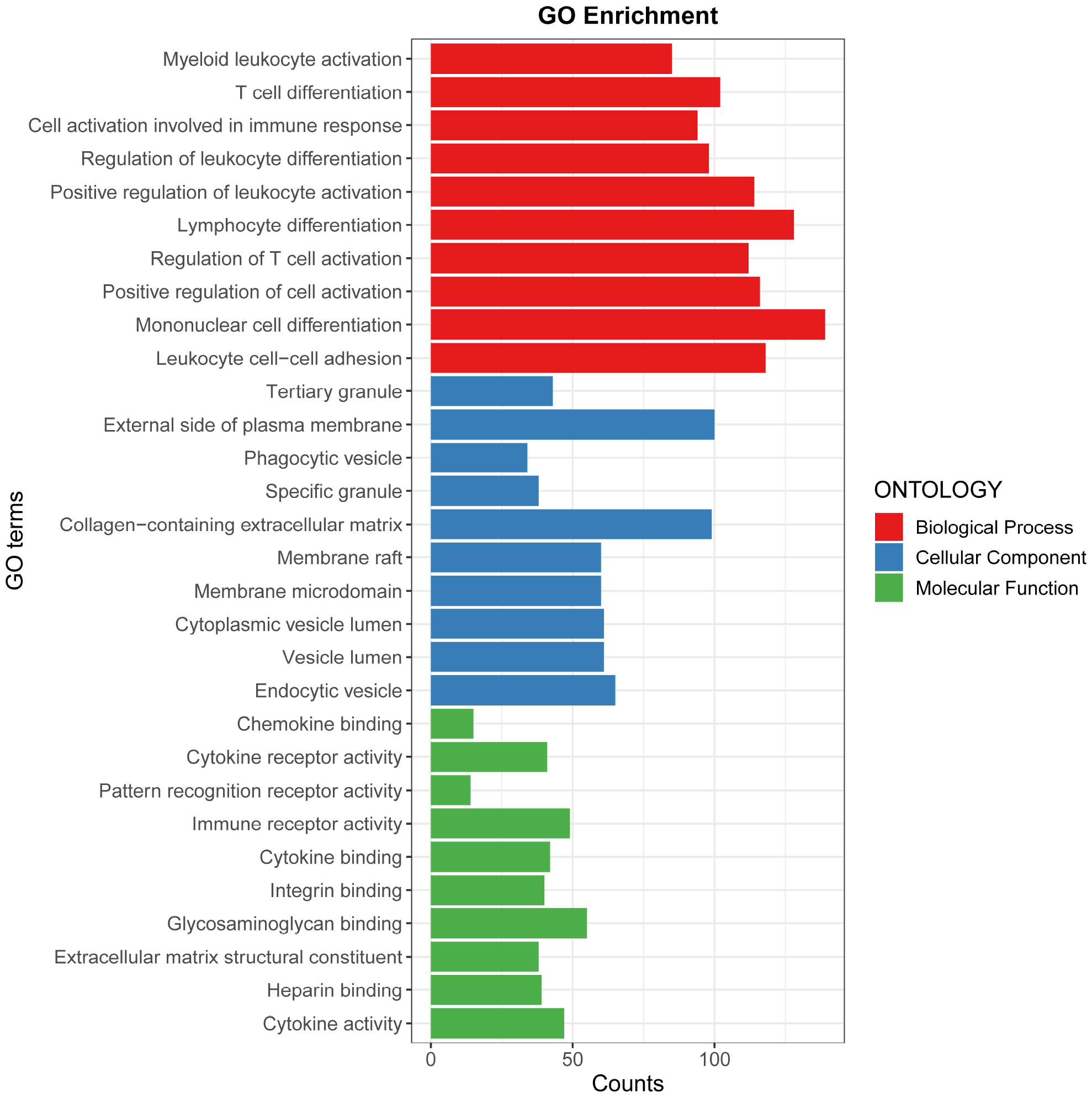
Top 10 results of Go enrichment analysis.

### Enrichment Analysis

Using the online tool String for KEGG enrichment analysis, the top 30 pathways identified were Cytokine-cytokine receptor interaction, PI3K-Akt signaling pathway, Neuroactive ligand-receptor interaction, MAPK signaling pathway, Chemokine signaling pathway, Cell adhesion molecules, Ras signaling pathway, Osteoclast differentiation, TNF signaling pathway, JAK-STAT signaling pathway, Focal adhesion, C-type lectin receptor signaling pathway, Leukocyte transendothelial migration, Calcium signaling pathway, FoxO signaling pathway, T cell receptor signaling pathway, NF-kappa B signaling pathway, Toll-like receptor signaling pathway, Apoptosis, NOD-like receptor signaling pathway, Phospholipase D signaling pathway, Wnt signaling pathway, Complement and coagulation cascades, Phagosome, ECM-receptor interaction, Natural killer cell mediated cytotoxicity, Platelet activation, Relaxin signaling pathway, Cell cycle, IL-17 signaling pathway. (Figure 2 and Table 1)

**Fig. 2.**
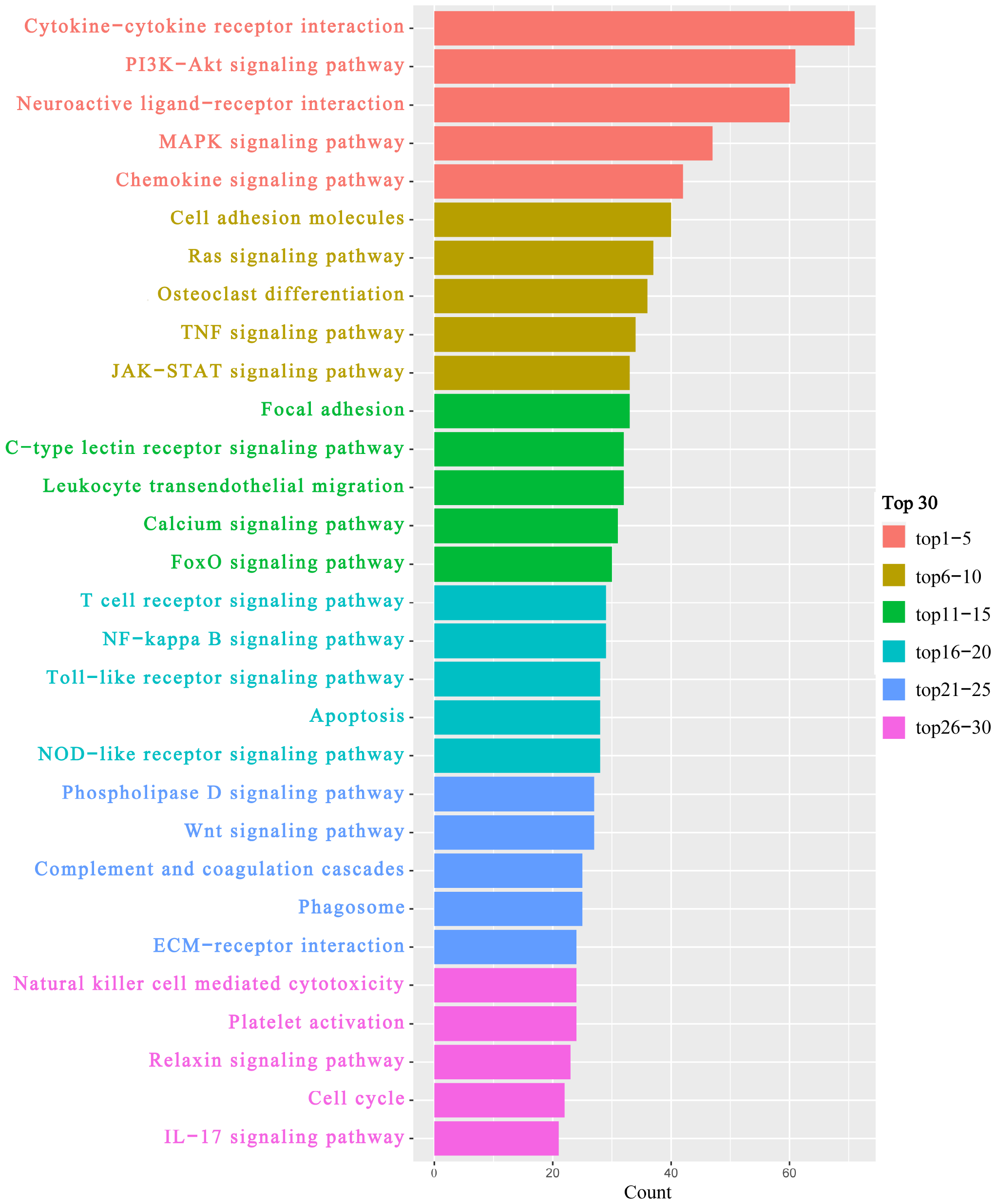
Top 30 results of KEGG enrichment analysis.

**Table 1.** Results of all KEGG enrichment assays after screening.

| #term ID | Description | Count | background<br>gene count | strength | false<br>discovery<br>rate |
| --- | --- | --- | --- | --- | --- |
| hsa04060 | Cytokine-cytokine receptor interaction | 71 | 282 | 0.42 | 5.67E-09 |
| hsa04151 | PI3K-Akt signaling pathway | 61 | 349 | 0.26 | 0.0004 |
| hsa04080 | Neuroactive ligand-receptor interaction | 60 | 329 | 0.28 | 0.00017 |
| hsa04010 | MAPK signaling pathway | 47 | 286 | 0.24 | 0.0055 |
| hsa04062 | Chemokine signaling pathway | 42 | 186 | 0.37 | 5.70E-05 |
| hsa04514 | Cell adhesion molecules | 40 | 138 | 0.48 | 3.04E-06 |
| hsa04014 | Ras signaling pathway | 37 | 225 | 0.24 | 0.0163 |
| hsa04380 | Osteoclast differentiation | 36 | 120 | 0.5 | 5.01E-06 |
| hsa04668 | TNF signaling pathway | 34 | 111 | 0.51 | 5.82E-06 |
| hsa04630 | JAK-STAT signaling pathway | 33 | 158 | 0.34 | 0.0014 |
| hsa04510 | Focal adhesion | 33 | 195 | 0.25 | 0.0177 |
| hsa04625 | C-type lectin receptor signaling pathway | 32 | 101 | 0.52 | 5.91E-06 |
| hsa04670 | Leukocyte transendothelial migration | 32 | 111 | 0.48 | 1.93E-05 |
| hsa04020 | Calcium signaling pathway | 31 | 191 | 0.23 | 0.0323 |
| hsa04068 | FoxO signaling pathway | 30 | 126 | 0.4 | 0.00042 |
| hsa04660 | T cell receptor signaling pathway | 29 | 100 | 0.48 | 4.80E-05 |
| hsa04064 | NF-kappa B signaling pathway | 29 | 101 | 0.48 | 5.19E-05 |
| hsa04620 | Toll-like receptor signaling pathway | 28 | 100 | 0.47 | 8.22E-05 |
| hsa04210 | Apoptosis | 28 | 131 | 0.35 | 0.0028 |
| hsa04621 | NOD-like receptor signaling pathway | 28 | 173 | 0.23 | 0.0422 |
| hsa04072 | Phospholipase D signaling pathway | 27 | 147 | 0.29 | 0.0163 |
| hsa04310 | Wnt signaling pathway | 27 | 154 | 0.26 | 0.0236 |
| hsa04610 | Complement and coagulation cascades | 25 | 82 | 0.51 | 8.14E-05 |
| hsa04145 | Phagosome | 25 | 141 | 0.27 | 0.028 |
| hsa04512 | ECM-receptor interaction | 24 | 88 | 0.46 | 0.00042 |
| hsa04650 | Natural killer cell mediated cytotoxicity | 24 | 120 | 0.32 | 0.0115 |
| hsa04611 | Platelet activation | 24 | 122 | 0.31 | 0.0133 |
| hsa04926 | Relaxin signaling pathway | 23 | 126 | 0.28 | 0.0281 |
| hsa04110 | Cell cycle | 22 | 120 | 0.28 | 0.0315 |
| hsa04657 | IL-17 signaling pathway | 21 | 91 | 0.38 | 0.0053 |
| hsa04725 | Cholinergic synapse | 20 | 109 | 0.28 | 0.0388 |
| hsa04666 | Fc gamma R-mediated phagocytosis | 19 | 90 | 0.35 | 0.0177 |
| hsa04659 | Th17 cell differentiation | 19 | 99 | 0.3 | 0.0335 |
| hsa04713 | Circadian entrainment | 18 | 91 | 0.32 | 0.0324 |
| hsa04750 | Inflammatory mediator regulation of TRP channels | 18 | 92 | 0.31 | 0.0336 |
| hsa04664 | Fc epsilon RI signaling pathway | 17 | 65 | 0.44 | 0.0053 |
| hsa04115 | p53 signaling pathway | 17 | 72 | 0.39 | 0.0119 |
| hsa04662 | B cell receptor signaling pathway | 17 | 78 | 0.36 | 0.0203 |
| hsa04658 | Th1 and Th2 cell differentiation | 17 | 85 | 0.32 | 0.0336 |
| hsa05340 | Primary immunodeficiency | 15 | 37 | 0.63 | 0.0004 |
| hsa02010 | ABC transporters | 12 | 45 | 0.45 | 0.0203 |

### Differential Gene Network Interaction Analysis

The top 50 core genes identified by Cytoscape software analysis of the STRING differential gene network interaction results were BUB1, CDK1, KIF11, CCNA2, KIF2C, CCNB1, TTK, BUB1B, DLGAP5, KIF20A, CDCA8, ASPM, TOP2A, BIRC5, NUSAP1, CENPF, CENPA, HJURP, CEP55, AURKB, CCNB2, PBK, MELK, HMMR, UBE2C, CDCA3, MKI67, KIF15, NEK2, CENPE, NUF2, SPAG5, KIF23, FOXM1, CDC45, CDKN3, PCLAF, RRM2, CDCA5, ESPL1, KIF14, DEPDC1, KIFC1, ANLN, ECT2, NCAPH, CDCA2, SKA1, SPC25, and OIP5. These genes were identified as playing key roles in the interactions within the differential gene network (Figure 3).

**Fig. 3.**
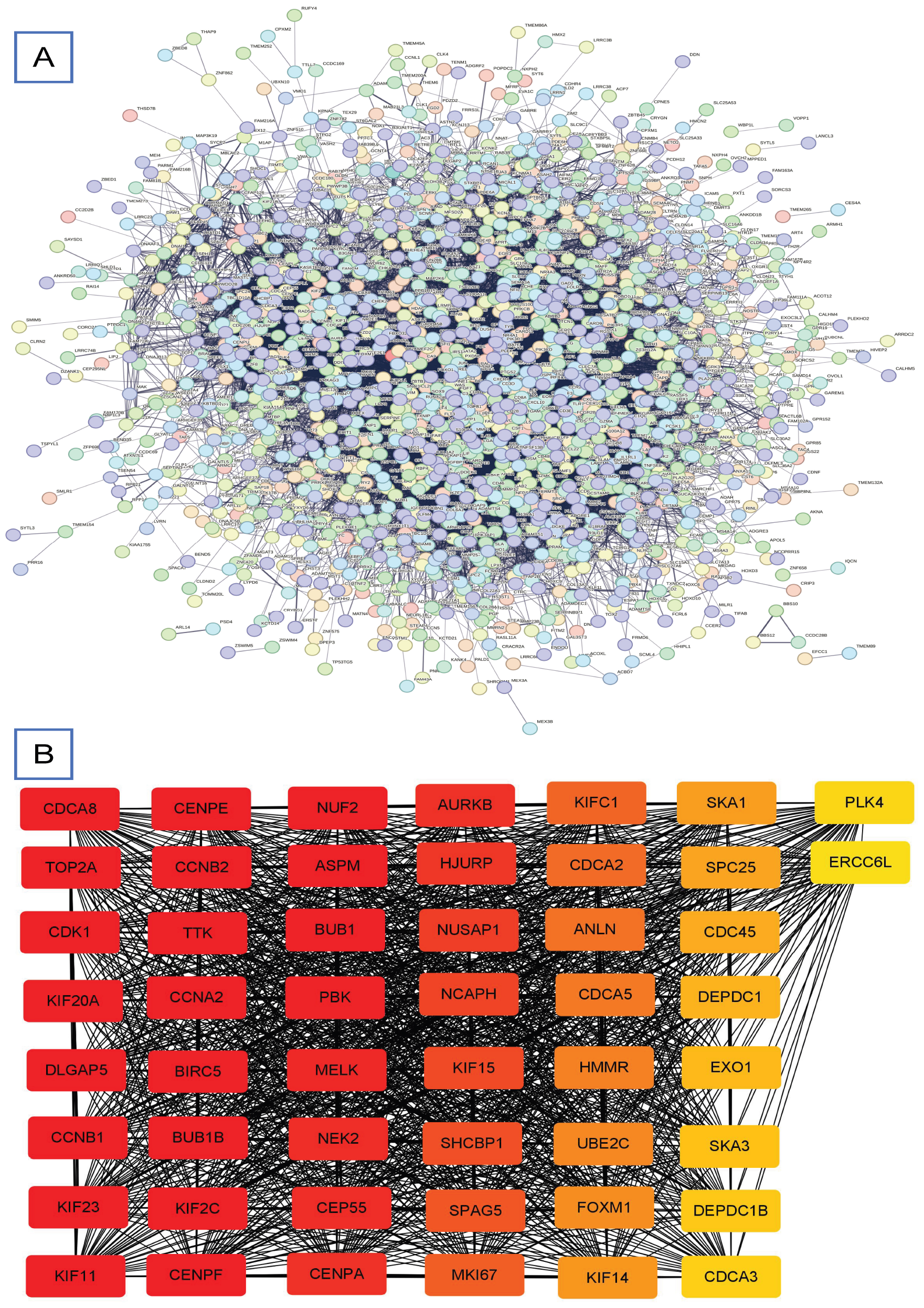
Protein-Protein Interaction (PPI) Network Analysis results.

## Discussion

The prevalence of calcium oxalate kidney stones within the kidney underlines the significance of this urological condition. To gain a deeper understanding of its pathogenesis and explore potential treatment methods, the establishment of a tree shrew calcium oxalate kidney stone model and gene transcriptome sequencing hold great promise.The selection of tree shrews, primates with a urinary system similar to humans, offers an experimental advantage in simulating human disease states. This model’s establishment facilitates a comprehensive understanding of calcium oxalate kidney stone formation and its associated pathophysiological alterations in tree shrews. This understanding not only reveals the underlying disease mechanisms but also serves as a valuable reference for human research. Complementing this, gene transcriptome sequencing provides an in-depth exploration of the molecular intricacies involved. This high-throughput sequencing technology allows for the identification of all transcripts within specific tissues or cells, offering insights into gene expression patterns and levels. By applying this sequencing approach to tree shrew kidney tissue, we can identify differentially expressed genes linked to calcium oxalate kidney stones. This, in turn, aids in the analysis of gene functions, interaction networks, and disease pathogenesis, potentially leading to the discovery of therapeutic targets.

In this study, we delve deeper into the significance of differential gene network interactions and their biological implications. Through the integration of the STRING database and Cytoscape software, we successfully identified 50 core genes playing pivotal roles in cellular processes such as the cell cycle, mitosis, and chromosome segregation. Genes like BUB1, CDK1, and KIF11 occupy central positions within these networks, regulating the cell cycle to ensure DNA replication and chromosomal segregation accuracy. KIF11, a member of the myosin family, is involved in mitotic spindle formation and chromosome movement. Its dysfunction can lead to chromosomal instability and uncontrolled cell proliferation. Additionally, genes closely associated with mitosis, such as CCNA2, KIF2C, and CCNB1, regulate processes like chromosome segregation, spindle assembly, and cell cycle checkpoints, vital for cell division’s smooth progression. Abnormalities in these genes can trigger mitotic errors, potentially leading to a range of cellular biological issues.

Biological process enrichment analysis underscores the importance of immune system-related processes. This alignment with existing medical knowledge underscores the immune system’s crucial role in various diseases, although its specific role in kidney stone formation remains incompletely understood. One plausible explanation is that the immune system modulates the kidney’s microenvironment, influencing processes like inflammatory responses and cytokine secretion, thereby affecting stone formation and progression. The KEGG enrichment analysis identifies potential therapeutic targets like cytokine-receptor interaction and PI3K-Akt signaling pathways. Drugs targeting key molecules within these pathways could potentially influence kidney stone formation and progression. Furthermore, these pathways could serve as diagnostic markers for early detection and monitoring of kidney stones.

In conclusion, our study successfully identified a set of core genes crucial for biological processes like the cell cycle, mitosis, and chromosome segregation. Abnormalities in these genes are closely linked to the occurrence and development of various diseases, highlighting the importance of further studying their

## Declarations

### Availability of data

The data is only used for learning and communication, please contact the corresponding author if required.

### Ethics statement

This study was reviewed and approved by the Animal Experiment Ethics Committee of Kunming Medical University (Approval no: kmmu20230176).

## References

[1] Tundo G, Vollstedt A, Meeks W, Pais V. Beyond Prevalence: Annual Cumulative Incidence of Kidney Stones in the United States. J Urol. 2021;205(6):1704–1709. doi:10.1097/JU.0000000000001629

[2] Huang Z, Wang G, Yang B, et al. Mechanism of ketotifen fumarate inhibiting renal calcium oxalate stone formation in SD rats. Biomed Pharmacother. 2022;151:113147. doi:10.1016/j.biopha.2022.113147

[3] Khan SR, Pearle MS, Robertson WG, et al. Kidney stones. Nat Rev Dis Primers. 2016;2:16008. Published 2016 Feb 25. doi:10.1038/nrdp.2016.8

[4] Fan Y, Huang ZY, Cao CC, et al. Genome of the Chinese tree shrew. Nat Commun. 2013. 4: 1426.

[5] Zhang J, Xiao H, Bi Y, et al. Characteristics of the tree shrew humoral immune system. Mol Immunol. 2020. 127: 175–185.

[6] Gu W, Li W, Wang W, et al. Response of the gut microbiota during the Clostridioides difficile infection in tree shrews mimics those in humans. BMC Microbiol. 2020. 20(1): 260.

[7] Chen Q, Ma ZX, Xia LB, et al. A tree shrew model for steroid-associated osteonecrosis. Zool Res. 2020. 41(5): 564–568.

[8] Wang G, Huang Z, Wu Y, Li P, Li J. Primate-like tree shrew will be the preferred animal for investigating nephrolithiasis. Urolithiasis. 2023;52(1):3. Published 2023 Nov 16. doi:10.1007/s00240-023-01507-6

