## Supplementary Table 1 for "Transcriptome sequencing and bioinformatics analysis of kidney tissue in a tree shrew model of calcium oxalate nephrolithiasis"

| Gene | logFC | P.Val |
| --- | --- | --- |
| MARCH1 | 2.64681025 | 2.30E-06 |
| A2M | 2.962505 | 4.76E-05 |
| A4GNT | 3.24108525 | 0.003284576 |
| AATK | 2.16647075 | 0.043690639 |
| ABCA1 | 1.78471775 | 0.00064759 |
| ABCA7 | 1.027568 | 0.032319122 |
| ABCA9 | 2.02923525 | 8.63E-05 |
| ABCB11 | -1.286993 | 0.001446826 |
| ABCC6 | -1.06638425 | 0.001737423 |
| ABCD2 | -2.8526605 | 0.023121483 |
| ABCG1 | 1.100039 | 0.025508158 |
| ABCG4 | 2.26799575 | 0.000384406 |
| ABCG5 | -2.21556975 | 0.022755629 |
| ABCG8 | -5.07160475 | 9.05E-05 |
| ABHD8 | -1.124579 | 0.00077255 |
| ABL2 | 1.03774275 | 0.000657662 |
| ACAP1 | 2.39276475 | 5.77E-06 |
| ACBD7 | 1.282202 | 0.037703092 |
| ACER2 | 1.20592875 | 0.000142833 |
| ACHE | 1.11666425 | 0.005074579 |
| ACOT12 | -1.4390835 | 0.023184904 |
| ACOXL | 2.12229475 | 0.027696279 |
| ACP4 | 2.51799975 | 0.049118151 |
| ACP7 | 3.90377325 | 0.001998902 |
| ACSBG1 | 1.31947025 | 9.40E-05 |
| ACSL5 | 1.21546875 | 0.030374174 |
| ACTL6B | 1.99432 | 0.033399123 |
| ACTN1 | 1.67662 | 9.57E-06 |
| ACVR1C | -1.552389 | 2.89E-05 |
| ADA | 1.6139265 | 0.001109148 |
| ADA2 | 2.49324475 | 0.000249864 |
| ADAM19 | 2.551076 | 0.001344389 |
| ADAM22 | -1.2338615 | 0.007124762 |
| ADAM28 | 3.7289615 | 0.000246443 |
| ADAM33 | 1.12953875 | 0.024720952 |
| ADAM8 | 1.6565575 | 0.000735327 |
| ADAMDEC1 | 4.32694525 | 0.003016221 |
| ADAMTS1 | 1.7081665 | 0.000205101 |
| ADAMTS14 | 2.7418065 | 0.003201744 |
| ADAMTS3 | 2.0858445 | 1.80E-05 |
| ADAMTS4 | 3.9071435 | 0.009826339 |
| ADAMTS5 | -1.00003525 | 0.006824638 |
| ADAMTS8 | 2.203154 | 0.001048605 |
| ADAMTSL1 | -1.3955195 | 0.000106234 |
| ADAMTSL3 | 1.017763 | 0.001641239 |
| ADCY7 | 2.092195 | 0.000139461 |
| ADCY8 | 1.2235685 | 0.034330974 |
| ADGRE1 | 4.14921425 | 0.002124346 |
| ADGRE3 | 3.90075075 | 0.001844597 |
| ADGRF2 | -2.424576 | 0.026666016 |
| ADGRG3 | 2.06408475 | 0.004379911 |
| ADGRG5 | 1.08642475 | 0.027607695 |
| ADGRG7 | -5.4506465 | 0.000469666 |
| ADGRV1 | -1.670809 | 0.004505666 |
| ADRA1B | -1.26503825 | 0.000365559 |
| ADRA1D | -3.134668 | 0.001628452 |
| ADRA2B | -1.05243175 | 0.031775731 |
| ADRB2 | 1.7709545 | 5.29E-05 |
| AEBP1 | 1.0067465 | 0.000206645 |
| AFF1 | 1.3375525 | 0.000257272 |
| AGAP2 | 2.07487375 | 0.002621086 |
| AGR2 | 1.49424975 | 0.000932887 |
| AGR3 | 2.09257275 | 0.046377846 |
| AHSG | 1.2805605 | 0.029853389 |
| AIF1 | 2.1360255 | 0.000152066 |
| AIFM2 | 2.27164925 | 9.67E-05 |
| AJUBA | -1.413506 | 2.08E-05 |
| AKAP12 | 3.40847225 | 0.000161704 |
| AKNA | 2.3614365 | 0.000265069 |
| ALAD | -1.00802125 | 0.000308058 |
| ALDH1A3 | 1.45630025 | 0.001310079 |
| ALDH3B2 | 1.71291925 | 0.001920049 |
| ALDH8A1 | -1.33783725 | 6.33E-05 |
| ALDOC | 1.51353925 | 0.045240404 |
| ALOX5AP | 1.7961895 | 1.13E-05 |
| ALPK1 | 3.462835 | 0.007147676 |
| AMBP | -1.0966245 | 0.000307972 |
| AMDHD1 | -1.147445 | 0.000165269 |
| AMER1 | -1.61471425 | 0.016228581 |
| AMOTL2 | -1.150452 | 0.044270545 |
| ANGPT2 | -1.417175 | 0.000789016 |
| ANGPTL4 | 4.818777 | 4.61E-06 |
| ANKDD1B | 2.56754875 | 0.039378875 |
| ANKRD34C | 1.93872225 | 0.011873962 |
| ANKRD50 | -1.07873875 | 0.000135494 |
| ANKRD61 | -1.41839525 | 0.000316169 |
| ANLN | 2.7060115 | 0.001365279 |
| ANXA1 | 1.051212 | 0.002679963 |
| ANXA13 | 2.67811725 | 7.67E-06 |
| ANXA2 | 1.18215975 | 0.000110851 |
| ANXA3 | 1.2448395 | 6.45E-05 |
| AOAH | 1.807831 | 0.000631044 |
| AP3B2 | 5.9597615 | 6.96E-08 |
| APBB1IP | 2.35986375 | 9.53E-05 |
| APCDD1L | -1.0543185 | 0.008826948 |
| APELA | -1.56025925 | 0.014322464 |
| APOA4 | 2.175016 | 0.032588143 |
| APOBEC3H | 1.05196975 | 0.034380076 |
| APOBR | 2.42356125 | 0.000254626 |
| APOH | -1.03071675 | 0.003861026 |
| APOL5 | -1.0942875 | 0.025356555 |
| APOLD1 | 3.32468675 | 0.000164267 |
| APRT | 1.0039955 | 0.00089644 |
| AQP10 | 2.31736525 | 0.01004286 |
| AQP9 | 4.96837725 | 0.000327596 |
| ARC | 2.72672275 | 0.000162773 |
| ARHGAP11A | 2.1234245 | 0.0013699 |
| ARHGAP15 | 1.06775625 | 0.010861296 |
| ARHGAP25 | 1.76317425 | 0.000712177 |
| ARHGAP26 | 1.340515 | 0.010857121 |
| ARHGAP30 | 2.6008455 | 5.05E-05 |
| ARHGAP36 | -3.9985885 | 0.0001433 |
| ARHGAP4 | 2.59073725 | 2.86E-05 |
| ARHGAP45 | 1.21780325 | 0.000570423 |
| ARHGAP9 | 1.6081845 | 0.001011352 |
| ARHGDIG | 2.422587 | 0.009730453 |
| ARHGEF37 | 1.3895385 | 0.000139496 |
| ARHGEF39 | 3.8593875 | 0.002802369 |
| ARID3C | 1.580482 | 0.012973048 |
| ARL11 | 4.37707175 | 0.001301097 |
| ARL14 | -1.04822275 | 0.001149465 |
| ARL4D | 1.36370625 | 0.008762465 |
| ARMC12 | 3.1597165 | 1.40E-05 |
| ARMC2 | 1.47875125 | 0.005334921 |
| ARMC3 | 4.0231895 | 7.18E-06 |
| ARMH1 | -1.214891 | 0.008551115 |
| ARNTL | 2.17762 | 0.001231152 |
| ARPC1B | 1.5715205 | 2.26E-05 |
| ARRB2 | 1.34801375 | 0.000194691 |
| ARRDC2 | 1.96297175 | 0.010484694 |
| ARSI | 2.715376 | 0.028105062 |
| ART4 | 3.66710875 | 0.000400068 |
| ASAH2 | 1.52263875 | 0.011206429 |
| ASB14 | -1.453728 | 0.02272833 |
| ASB2 | -1.258976 | 0.036321963 |
| ASB5 | 2.9710425 | 0.004542972 |
| ASCL2 | 1.18424125 | 0.034374194 |
| ASCL4 | -1.30084725 | 0.00087114 |
| ASF1B | 1.6090015 | 0.003443946 |
| ASPA | -1.18175625 | 0.000575583 |
| ASPM | 2.20984825 | 0.007354716 |
| ASPN | 1.614704 | 0.01031111 |
| ASTN2 | 1.090598 | 0.020433884 |
| ASZ1 | 4.4477655 | 2.59E-06 |
| ATCAY | 2.19615875 | 0.03515431 |
| ATF3 | 3.87001975 | 0.000397985 |
| ATOH7 | -1.15044025 | 0.02843496 |
| ATP12A | -1.52835275 | 0.00159076 |
| ATP8B4 | 2.496677 | 0.000360911 |
| ATXN7L1 | 1.389603 | 0.001390168 |
| AURKB | 1.30781075 | 0.00353233 |
| AVPR1A | -1.166756 | 0.000619438 |
| B2M | 1.03696625 | 0.002771901 |
| B3GALT1 | -2.72672275 | 0.001090054 |
| B3GNT3 | 2.98323825 | 0.01299773 |
| BAALC | 2.9534455 | 1.61E-06 |
| BAMBI | -1.15604075 | 3.37E-05 |
| BANK1 | 1.453234 | 0.007336573 |
| BASP1 | 1.91210825 | 0.000190822 |
| BATF | 1.16988175 | 0.011183272 |
| BATF2 | 1.24104525 | 0.006946595 |
| BBS10 | -1.18283375 | 0.007269112 |
| BBS12 | -1.1012315 | 0.001668566 |
| BCAS1 | -1.4162735 | 0.049033139 |
| BCAT1 | 1.10188475 | 0.031115673 |
| BCL11B | 1.63370375 | 0.000811231 |
| BCL2L11 | 1.37734275 | 0.000173557 |
| BCL3 | 1.1775175 | 0.015114074 |
| BCL6 | 1.72500525 | 0.002232296 |
| BCL7A | -1.3339525 | 0.001690494 |
| BDKRB1 | 2.6434245 | 0.000985609 |
| BDKRB2 | 3.192846 | 1.90E-06 |
| BEND3 | -1.18060625 | 0.000553336 |
| BEND5 | -1.151614 | 3.15E-05 |
| BEST1 | 2.8115585 | 0.000126452 |
| BEST4 | 1.49432025 | 0.027862782 |
| BFSP2 | 1.191217 | 0.001561567 |
| BGN | 1.1827205 | 0.002592761 |
| BHLHA15 | 1.2005755 | 0.04943691 |
| BHLHE41 | 1.1963 | 0.003146838 |
| BHMG1 | 2.22672275 | 7.99E-05 |
| BID | 1.524511 | 0.000515959 |
| BIN2 | 2.7981465 | 2.60E-05 |
| BIRC2 | 1.60215175 | 1.26E-05 |
| BIRC3 | 1.96671375 | 0.001831656 |
| BIRC5 | 3.4554455 | 0.002568275 |
| BLK | 1.50904325 | 0.040959014 |
| BLM | 2.04291225 | 0.000642464 |
| BMF | 1.9342735 | 3.94E-05 |
| BMPER | 2.14598425 | 0.002032763 |
| BRAP | -1.1781725 | 0.000189393 |
| BRCA2 | 1.454519 | 0.00296142 |
| BRINP3 | 1.02917075 | 0.009319773 |
| BRIP1 | 1.63136275 | 0.010894051 |
| BRSK2 | 2.394829 | 0.0175107 |
| BTBD11 | 2.1689855 | 0.000661993 |
| BTG1 | 1.03680225 | 0.000875126 |
| BTG2 | 1.598177 | 0.002580761 |
| BTK | 2.2690455 | 0.00043001 |
| BUB1 | 4.2534625 | 0.003141779 |
| BUB1B | 2.44235 | 0.00065128 |
| C1QA | 1.76726475 | 0.000367794 |
| C1QB | 1.715009 | 9.45E-05 |
| C1QC | 1.851325 | 0.000338513 |
| C1QTNF2 | -1.00644225 | 0.001533068 |
| C3 | 2.27713775 | 0.000305285 |
| C4BPA | 3.24668675 | 9.56E-07 |
| C5 | 1.734506 | 0.001517453 |
| C5AR1 | 2.6938465 | 0.00014861 |
| CA13 | 1.8089705 | 0.005958692 |
| CA9 | 1.31177875 | 0.000142364 |
| CACNA1B | 3.333678 | 0.00028456 |
| CACNA1I | 3.23772825 | 0.034117418 |
| CACNG5 | 2.34158075 | 0.006262394 |
| CALCR | -2.434728 | 0.003330461 |
| CALHM4 | -1.503153 | 0.003408653 |
| CALHM5 | -1.34566 | 0.000994254 |
| CALML6 | -1.184573 | 0.016149382 |
| CAMK2B | 1.26992525 | 0.005800799 |
| CAMK4 | 1.3443595 | 0.00036766 |
| CAPG | 3.54461 | 3.37E-05 |
| CAPN9 | -1.84247 | 0.007747324 |
| CARD11 | 2.652872 | 0.000330955 |
| CARD14 | 2.13813525 | 0.012805955 |
| CARD8 | 4.0186265 | 3.69E-05 |
| CARD9 | 1.9264135 | 0.019118526 |
| CARMIL2 | 2.5203585 | 0.00034719 |
| CASC1 | 1.14004575 | 0.012330034 |
| CASS4 | -1.67624725 | 0.000529143 |
| CAVIN2 | -1.33921225 | 0.000133137 |
| CBFA2T3 | 1.42519875 | 0.01618022 |
| CBLB | 1.5668235 | 0.000207047 |
| CBR3 | 1.02325025 | 0.002890013 |
| CBX8 | -1.34036525 | 0.004192796 |
| CC2D2B | -1.1728065 | 0.000425242 |
| CCDC102A | -1.06841925 | 0.011850424 |
| CCDC114 | 1.0942395 | 0.001115359 |
| CCDC146 | -1.37148 | 0.009666554 |
| CCDC162P | 2.11860925 | 0.010854766 |
| CCDC169 | 2.336074 | 0.010594812 |
| CCDC180 | 2.46527075 | 0.000177948 |
| CCDC197 | -1.3198415 | 0.038593502 |
| CCDC28B | 1.4571355 | 0.010604069 |
| CCDC40 | 1.508466 | 0.044838305 |
| CCDC69 | 1.16363175 | 0.001030243 |
| CCDC80 | 2.605634 | 8.34E-05 |
| CCER2 | 1.254387 | 0.002823227 |
| CCL14 | 2.271379 | 0.000165672 |
| CCL15 | 2.87317025 | 1.43E-06 |
| CCL19 | 4.31362175 | 0.005704236 |
| CCL22 | 7.17905325 | 6.49E-06 |
| CCL24 | 5.22569 | 3.74E-05 |
| CCL27 | -1.15949925 | 0.021660546 |
| CCL5 | 2.91421175 | 1.94E-05 |
| CCN2 | 1.959601 | 0.000343819 |
| CCN5 | 3.37909875 | 8.63E-07 |
| CCNA2 | 1.47712475 | 0.017240115 |
| CCNB1 | 1.83862325 | 0.005456732 |
| CCNB2 | 4.16342925 | 0.00177455 |
| CCNF | 1.34840675 | 0.044127534 |
| CCNL1 | 1.591515 | 0.000510021 |
| CCR2 | 1.437246 | 0.017451138 |
| CCR6 | 3.16166775 | 0.005128532 |
| CCR8 | 3.14454625 | 0.001637616 |
| CCRL2 | 2.451427 | 0.036361372 |
| CD109 | 4.037187 | 5.42E-06 |
| CD14 | 2.76475 | 0.000138658 |
| CD163 | 4.04597 | 5.64E-07 |
| CD177 | 2.11285725 | 0.012755727 |
| CD180 | 1.36158475 | 0.021767762 |
| CD2 | 2.7624245 | 5.14E-05 |
| CD207 | 3.89185275 | 0.000863168 |
| CD226 | 1.61227975 | 0.003731593 |
| CD244 | -1.62266725 | 0.002041567 |
| CD27 | 4.3097005 | 0.001431655 |
| CD274 | 1.873394 | 0.000356489 |
| CD28 | 4.71029525 | 0.001803064 |
| CD37 | 2.638147 | 0.000242573 |
| CD3D | 2.22476375 | 0.003947561 |
| CD3E | 3.02066925 | 1.44E-06 |
| CD3G | 1.92694275 | 0.003144953 |
| CD4 | 3.2558465 | 0.000235514 |
| CD44 | 2.59803925 | 4.36E-07 |
| CD46 | 1.42835525 | 0.007277597 |
| CD48 | 3.8496805 | 2.03E-06 |
| CD5 | 3.78687075 | 3.83E-06 |
| CD53 | 2.54184425 | 2.23E-05 |
| CD5L | 1.8072045 | 0.033015853 |
| CD6 | 3.59176125 | 9.38E-05 |
| CD68 | 2.69782825 | 2.33E-05 |
| CD69 | 2.83874925 | 0.005350368 |
| CD70 | 2.52713125 | 0.018742308 |
| CD72 | 4.1615245 | 3.18E-06 |
| CD74 | 2.0936305 | 1.53E-05 |
| CD83 | 3.66109 | 2.67E-05 |
| CD84 | 5.07460275 | 1.35E-05 |
| CD86 | -1.00997 | 0.001090081 |
| CD8A | 5.09693775 | 0.000854215 |
| CD8B | 3.223199 | 0.009180481 |
| CD93 | 1.38948325 | 0.00945254 |
| CD96 | 2.9966635 | 0.000149237 |
| CD99 | 1.259445 | 0.000228564 |
| CDC20B | 2.363832 | 0.01387317 |
| CDC25B | 1.1259745 | 0.005627846 |
| CDC42EP2 | 1.5196245 | 0.001541473 |
| CDC42EP4 | 1.07363675 | 0.000722312 |
| CDC45 | 3.0329645 | 0.028407037 |
| CDC6 | 2.006708 | 0.001291216 |
| CDCA2 | 1.0234125 | 0.008610044 |
| CDCA3 | 3.8589265 | 0.014269867 |
| CDCA5 | 1.7133935 | 0.001594825 |
| CDCA7 | -1.483713 | 0.028132943 |
| CDCA8 | 1.31313975 | 0.005075427 |
| CDCP1 | 1.4689515 | 0.00060454 |
| CDCP2 | 3.4774265 | 1.51E-06 |
| CDH20 | -1.3369655 | 0.016344374 |
| CDH23 | 1.90585325 | 1.68E-06 |
| CDHR4 | 2.34158075 | 0.015332692 |
| CDK1 | 1.26137975 | 0.001368924 |
| CDKN1A | 3.02217525 | 2.11E-06 |
| CDKN3 | 1.376175 | 0.001223629 |
| CDNF | -1.68368075 | 0.00195208 |
| CDSN | 1.584627 | 0.043290917 |
| CDT1 | 1.23171025 | 0.046708825 |
| CEBPD | 1.33063825 | 0.001348692 |
| CELF5 | -2.59288975 | 0.012218378 |
| CELSR3 | 2.34158075 | 0.003233236 |
| CEMIP2 | 2.281997 | 3.05E-06 |
| CENPA | 2.206568 | 0.005742382 |
| CENPE | 1.992698 | 0.000833553 |
| CENPF | 3.197541 | 0.000374799 |
| CENPI | 1.267694 | 0.000664929 |
| CENPK | 2.11191975 | 0.007716659 |
| CENPS | 3.4159515 | 0.007209825 |
| CENPT | 1.9696805 | 0.004063716 |
| CENPU | 1.036527 | 0.016741843 |
| CENPW | 1.88033975 | 0.000858987 |
| CEP295NL | 2.0066125 | 0.007147206 |
| CEP55 | 2.5550975 | 0.012255877 |
| CER1 | 3.98820225 | 0.001088023 |
| CERCAM | 1.463919 | 0.002163134 |
| CERKL | 1.55235925 | 0.002056739 |
| CES4A | 2.21759125 | 0.018507502 |
| CFAP126 | 4.09195575 | 0.000694646 |
| CFAP45 | 2.7785885 | 0.00024276 |
| CFAP52 | 1.8100785 | 0.043573921 |
| CFAP54 | 1.44770375 | 0.010483958 |
| CFLAR | 2.271712 | 8.60E-07 |
| CFP | 2.3665425 | 0.00018719 |
| CFTR | 3.81346675 | 0.000173151 |
| CGAS | 2.60389025 | 0.000186066 |
| CH25H | 4.95201125 | 0.00119549 |
| CHEK2 | 1.036569 | 0.002598214 |
| CHI3L1 | 4.0318325 | 3.46E-06 |
| CHIT1 | 14.04555225 | 1.48E-06 |
| CHKA | 1.00475075 | 0.000644278 |
| CHL1 | 2.66943025 | 0.011151992 |
| CHP2 | -2.4181065 | 0.015952413 |
| CHRD | 3.3837655 | 0.003491725 |
| CHRM4 | -1.27906375 | 0.002682505 |
| CHRNA2 | -2.142464 | 0.027357022 |
| CHRNB1 | 1.301647 | 0.00084502 |
| CHST2 | 1.05302775 | 0.022789266 |
| CHST3 | -1.53458125 | 0.002592245 |
| CHST7 | -2.274436 | 0.032763169 |
| CIART | 2.00931175 | 0.004650451 |
| CIDEA | -1.93041575 | 0.006663792 |
| CIITA | 1.3176365 | 0.007643091 |
| CILP | 3.5822385 | 0.005051886 |
| CIP2A | 1.0921705 | 0.001905143 |
| CISH | 2.03553625 | 0.001256249 |
| CKAP2 | 2.36927325 | 0.001891048 |
| CKAP2L | 1.9438825 | 0.004537927 |
| CLCF1 | 1.54781675 | 0.005074268 |
| CLDN11 | 5.768761 | 0.002878702 |
| CLDN14 | -1.19197975 | 0.024032685 |
| CLDN17 | 1.410964 | 0.027217924 |
| CLDN23 | -1.59101475 | 0.003175194 |
| CLDN34 | 1.580482 | 0.012973048 |
| CLDN4 | 1.55046525 | 0.008846253 |
| CLDN9 | 2.05720475 | 0.014450958 |
| CLDND2 | 1.58894825 | 0.023435254 |
| CLEC4D | 4.89405625 | 0.000244978 |
| CLEC4E | 8.165172 | 5.30E-05 |
| CLEC5A | 1.68630225 | 0.013877901 |
| CLEC6A | 5.55447325 | 2.57E-09 |
| CLEC7A | 4.14143125 | 4.74E-07 |
| CLIC6 | -1.02782375 | 0.001183229 |
| CLK1 | -1.20827225 | 0.001401828 |
| CLK4 | 1.12612175 | 0.000954286 |
| CLNK | 4.302793 | 0.000239525 |
| CLP1 | -1.0033285 | 0.006289567 |
| CLRN2 | 1.35554175 | 0.000814316 |
| CLSPN | 2.92444 | 0.000298843 |
| CLTRN | -1.1709165 | 0.004261879 |
| CMKLR1 | 1.61448425 | 0.004446346 |
| CMTM5 | -1.246248 | 0.026526067 |
| CMTM6 | 1.82176875 | 0.034858343 |
| CNNM1 | 5.19594925 | 4.54E-05 |
| CNTLN | 3.58799625 | 4.54E-06 |
| CNTN5 | -1.85495975 | 0.000161562 |
| CNTNAP2 | -1.06181475 | 0.001612609 |
| COCH | -1.03709175 | 0.010330822 |
| COL13A1 | 1.2279615 | 0.001097608 |
| COL1A1 | 1.5851885 | 0.000348299 |
| COL1A2 | 1.22928775 | 0.005390232 |
| COL21A1 | -1.06319025 | 0.005521972 |
| COL22A1 | 1.18647625 | 0.046664456 |
| COL28A1 | 2.17511 | 0.021486157 |
| COL3A1 | 1.031873 | 0.006410322 |
| COL4A1 | 1.09100025 | 0.001232241 |
| COL5A2 | 1.047476 | 0.000864354 |
| COL6A1 | 1.0711345 | 0.003413242 |
| COL6A3 | 1.05627875 | 0.005396162 |
| COL7A1 | 2.27920425 | 0.043190491 |
| COMMD6 | -2.75577925 | 0.015880216 |
| CORO1A | 1.91251725 | 0.000296162 |
| CORO1C | 1.007875 | 0.000390758 |
| CORO2A | 1.6964645 | 0.002811372 |
| COTL1 | 1.26540325 | 0.000119869 |
| CPNE5 | 1.23153625 | 0.022710279 |
| CPVL | 1.3234975 | 0.005707151 |
| CPXM1 | 1.58358825 | 0.003683721 |
| CPXM2 | 1.36083 | 0.000171591 |
| CPZ | 2.12571775 | 0.00076704 |
| CRACR2A | 3.77441025 | 0.001452757 |
| CREB3L3 | -1.655011 | 0.008137261 |
| CRHBP | 2.0627405 | 0.026359864 |
| CRIP3 | -5.422408 | 1.07E-08 |
| CRISPLD2 | 2.5026515 | 0.001474995 |
| CRLF1 | 3.35967375 | 6.67E-05 |
| CRLF2 | 2.69854775 | 1.15E-05 |
| CRTAM | 2.6376865 | 0.001523507 |
| CRYBB3 | 2.3729635 | 0.002201435 |
| CRYBG1 | 1.27761975 | 0.000419634 |
| CRYBG2 | 2.28174775 | 0.034976332 |
| CRYGN | 3.324802 | 3.73E-05 |
| CSF1 | 1.33155 | 0.002915988 |
| CSF1R | 3.09479975 | 3.12E-06 |
| CSF2RA | 5.61536625 | 0.000118718 |
| CSF2RB | 1.7250105 | 0.000822275 |
| CSF3 | 1.84334725 | 0.03623195 |
| CSF3R | 2.57359775 | 0.000205192 |
| CSMD2 | -1.03251925 | 7.85E-05 |
| CSRNP1 | 2.4208915 | 0.001428185 |
| CST6 | 4.22700425 | 0.000130581 |
| CSTB | 1.39825575 | 0.001529002 |
| CTHRC1 | 4.654155 | 0.000310505 |
| CTLA4 | 4.770532 | 0.000243157 |
| CTNNA2 | -1.30282 | 0.04102966 |
| CTRC | -1.323225 | 0.044445426 |
| CTSC | 1.0015885 | 0.000258322 |
| CTSH | 2.778782 | 0.000175387 |
| CTSK | 1.98746525 | 0.000415517 |
| CTSS | 4.26054875 | 4.88E-06 |
| CTSW | 2.26777675 | 0.000181626 |
| CUNH10orf105 | 3.0231895 | 0.003047429 |
| CUNH10orf62 | -1.160964 | 0.021365242 |
| CUNH11orf86 | 1.25663525 | 0.047298566 |
| CUNH11orf88 | 2.19615875 | 0.012018667 |
| CUNH11orf96 | 2.3987935 | 0.00186195 |
| CUNH12orf43 | -1.00183225 | 0.000210208 |
| CUNH12orf60 | 3.90727525 | 0.002277865 |
| CUNH16orf54 | 2.4041015 | 0.000162152 |
| CUNH16orf82 | -2.5979 | 0.000229095 |
| CUNH17orf67 | 2.194213 | 0.002016581 |
| CUNH19orf38 | 2.34772775 | 0.037242852 |
| CUNH19orf44 | -1.86712675 | 0.016009719 |
| CUNH19orf67 | -1.0315395 | 0.015770043 |
| CUNH1orf158 | 3.99592675 | 7.76E-05 |
| CUNH1orf162 | 1.10589925 | 0.00131583 |
| CUNH20orf141 | 2.73231475 | 3.46E-05 |
| CUNH20orf204 | -1.3618255 | 0.047194737 |
| CUNH22orf31 | 3.05358025 | 5.07E-06 |
| CUNH2orf73 | -1.04526675 | 0.021779104 |
| CUNH2orf92 | 2.94032875 | 0.013735504 |
| CUNH3orf18 | -1.285382 | 6.93E-05 |
| CUNH3orf49 | 1.87296325 | 0.016503383 |
| CUNH4orf3 | -1.39624075 | 0.014101715 |
| CUNH4orf54 | 1.99136375 | 0.000834143 |
| CUNH5orf49 | 2.87694875 | 0.008379219 |
| CUNH5orf58 | 4.43575275 | 0.000340041 |
| CUNH6orf118 | 1.9285615 | 0.021830278 |
| CUNH7orf25 | 7.00691275 | 0.006580937 |
| CUNH8orf89 | 2.751757 | 3.60E-08 |
| CXCL10 | 2.89587175 | 2.22E-06 |
| CXCL11 | 2.4004965 | 1.78E-05 |
| CXCL14 | 2.9760825 | 0.033248595 |
| CXCL16 | 1.7280935 | 4.58E-05 |
| CXCL8 | 4.6248575 | 0.000772905 |
| CXCL9 | 3.087345 | 8.38E-06 |
| CXCR1 | 2.52634625 | 0.018774333 |
| CXCR2 | 3.23778575 | 0.000523738 |
| CXCR3 | 4.3353275 | 0.000186195 |
| CXCR4 | 1.22852025 | 0.000377919 |
| CXCR6 | 1.13015325 | 0.010156375 |
| CYSLTR2 | 1.6432835 | 0.004110203 |
| DACT2 | -1.76179575 | 3.45E-05 |
| DAPP1 | 1.3375735 | 0.003977619 |
| DAW1 | 3.8710935 | 0.000681767 |
| DAZAP2 | 1.333059 | 0.028176844 |
| DBF4 | 1.14450025 | 0.00837633 |
| DCSTAMP | 10.06112475 | 1.78E-06 |
| DDIT4 | 1.66404875 | 0.000620043 |
| DDIT4L | -3.7598665 | 1.25E-07 |
| DDN | -1.6579665 | 0.000207612 |
| DDX11 | 1.2349825 | 0.000435968 |
| DEF6 | 2.00104225 | 0.002338871 |
| DEFB134 | 2.05720475 | 0.014450958 |
| DEPDC1 | 3.625526 | 0.012425694 |
| DEPDC1B | 1.76786025 | 0.000819681 |
| DEPP1 | 2.0421045 | 0.001147103 |
| DEPTOR | -1.03284825 | 0.002529419 |
| DERL3 | 1.156219 | 0.017002155 |
| DGKE | 1.86927475 | 0.000378674 |
| DIAPH3 | 2.31469 | 0.001289018 |
| DIO3 | 3.7145985 | 0.010100827 |
| DKK1 | 5.14605575 | 0.002511284 |
| DKK3 | 1.30165125 | 0.000961423 |
| DLG2 | -1.4951925 | 0.015313753 |
| DLGAP2 | -2.30720475 | 0.012978852 |
| DLGAP5 | 2.475554 | 0.001075387 |
| DMP1 | 2.265367 | 0.023182657 |
| DMRT3 | 3.1229635 | 0.013510098 |
| DNAAF3 | 1.637376 | 0.000565963 |
| DNAH17 | 2.44738325 | 0.000168236 |
| DNAH2 | 1.5860465 | 0.018527262 |
| DNAH6 | 1.11270625 | 0.00153968 |
| DNAH7 | 2.2774575 | 0.012673586 |
| DNAH8 | -2.7259935 | 0.01086521 |
| DNAI1 | 2.3882425 | 0.032954728 |
| DNAJA4 | 1.6451845 | 0.014217828 |
| DNAJB13 | 3.8253895 | 0.000257821 |
| DNAJB2 | -1.127443 | 2.25E-05 |
| DNAJC5B | 1.7924815 | 0.013538637 |
| DNASE2 | 1.2138435 | 0.00350773 |
| DNMT3B | -1.52064625 | 0.000100176 |
| DOCK2 | 2.4300995 | 2.06E-05 |
| DOK2 | 2.06230925 | 0.001336237 |
| DOK5 | -1.3080055 | 0.030193208 |
| DPEP2 | 5.772466 | 3.14E-07 |
| DPEP3 | 2.824802 | 0.005057446 |
| DPF3 | -1.8251835 | 0.003168738 |
| DPPA3 | 2.49432 | 0.012665632 |
| DPYSL2 | 1.19060225 | 0.017953178 |
| DPYSL5 | 3.07694275 | 2.84E-05 |
| DRAXIN | 2.1785615 | 0.017504357 |
| DRC1 | 1.42114175 | 0.044327769 |
| DRC3 | 2.63686525 | 0.001445975 |
| DRC7 | 1.75732175 | 0.043662441 |
| DSCAML1 | 2.58863225 | 0.000120561 |
| DSPP | 2.7034455 | 3.26E-06 |
| DTD2 | -1.1042615 | 0.006083615 |
| DTX1 | 1.1178865 | 0.005999025 |
| DUOX2 | 3.3889245 | 0.00282093 |
| DUSP1 | 2.24218925 | 0.000291196 |
| DUSP10 | 1.94660575 | 0.014249358 |
| DUSP2 | 4.549011 | 0.003550094 |
| DUSP5 | 2.4398025 | 0.002250447 |
| DUSP6 | 1.32703775 | 0.004074735 |
| DUSP8 | 1.85018275 | 0.004992622 |
| DYDC2 | 4.146318 | 2.33E-08 |
| DZANK1 | 1.639582 | 0.000964442 |
| E2F1 | 2.6237495 | 5.17E-05 |
| E2F7 | 2.4200805 | 0.00215925 |
| E2F8 | 3.587449 | 0.003969629 |
| ECEL1 | 2.599686 | 0.012336111 |
| ECT2 | 2.35440775 | 0.000372122 |
| EDN2 | 3.365562 | 1.10E-05 |
| EFCC1 | 1.98142325 | 0.01046307 |
| EFEMP2 | 1.06937975 | 0.000677184 |
| EFHC1 | 2.047753 | 0.006203562 |
| EFNB3 | -1.527199 | 0.001405614 |
| EGFL6 | -1.06254325 | 0.041883814 |
| EGR1 | 2.43029525 | 0.005530841 |
| EGR2 | 2.97967 | 0.00023451 |
| EGR3 | 2.4626005 | 0.005995113 |
| ELAVL2 | -2.718552 | 0.000142657 |
| ELL | 1.77839925 | 0.000779987 |
| ELL2 | 2.62427475 | 5.77E-06 |
| EMB | 2.4178615 | 0.016589878 |
| EMILIN1 | 1.554064 | 0.000170373 |
| EMILIN3 | -3.351043 | 0.00561416 |
| EMP3 | 1.41998825 | 0.001395304 |
| ENDOU | 1.89709175 | 0.025015378 |
| ENOX1 | -1.1851745 | 0.001467987 |
| EOMES | 3.16065375 | 0.005706108 |
| EPHA10 | 1.41715075 | 0.018245982 |
| EPHA7 | -1.00626 | 0.010797735 |
| EPHB2 | 1.841805 | 0.004235074 |
| EPOP | 2.8149885 | 0.00012622 |
| EPYC | 4.33952525 | 0.013203748 |
| ERAS | -3.97104275 | 5.50E-08 |
| ERCC6L | 1.90737775 | 0.000776238 |
| ERMN | -3.7954475 | 0.048854112 |
| ERRFI1 | 3.28904725 | 6.52E-07 |
| ESCO2 | 2.210452 | 0.01378813 |
| ESM1 | -1.23954825 | 0.000199115 |
| ESPL1 | 2.76161375 | 0.00047672 |
| ESPN | -1.76649575 | 0.000523915 |
| ETFBKMT | -1.429759 | 0.006859129 |
| ETV4 | 2.48115625 | 0.021870484 |
| ETV5 | 1.79315825 | 5.45E-05 |
| ETV7 | 3.705118 | 0.003377699 |
| EVA1A | 1.72304875 | 3.79E-06 |
| EVA1C | 3.05459425 | 1.65E-05 |
| EVI2A | 2.07844875 | 0.000121454 |
| EVI2B | 2.04880975 | 0.00060218 |
| EXD1 | 1.0882065 | 0.001729766 |
| EXO1 | 1.3970325 | 0.009437924 |
| EXOC3L2 | -2.4753885 | 4.25E-05 |
| F13A1 | 1.5243 | 0.001392249 |
| F2 | 1.00172975 | 0.02148236 |
| F2RL2 | 1.2492415 | 0.001959904 |
| F3 | 1.43349425 | 0.000246235 |
| FABP1 | -1.16624525 | 0.003354652 |
| FAM102A | 1.13199825 | 0.002521353 |
| FAM111A | 2.063106 | 0.00342675 |
| FAM129A | 1.28169725 | 0.000299044 |
| FAM129B | 1.05121775 | 0.002926885 |
| FAM133A | -2.6376865 | 0.002397245 |
| FAM151B | -1.436844 | 0.003703474 |
| FAM155B | -1.3628285 | 0.010329067 |
| FAM162B | -4.54039325 | 0.000675618 |
| FAM163A | -2.50874925 | 0.039224696 |
| FAM169B | -1.4325795 | 0.000380794 |
| FAM170B | 1.4959855 | 0.017363756 |
| FAM183A | 3.32480225 | 2.41E-06 |
| FAM216A | 1.4581795 | 0.013494049 |
| FAM216B | 4.6542845 | 0.000172813 |
| FAM227A | -2.70631925 | 0.001446025 |
| FAM43A | 1.101664 | 0.004893242 |
| FAM43B | -1.68688275 | 0.000100491 |
| FAM50B | 3.384062 | 0.000517858 |
| FAM72A | 3.095147 | 0.014594869 |
| FAM81B | 3.16544475 | 4.30E-06 |
| FAM83A | 2.74432 | 0.014448474 |
| FAM83D | 2.20095725 | 0.000665403 |
| FAM83E | 3.2731895 | 4.64E-06 |
| FAM84A | -1.40718475 | 7.66E-05 |
| FAM98A | -1.34445025 | 6.92E-05 |
| FANCD2 | 2.08008375 | 4.74E-05 |
| FANCE | 1.8504135 | 0.003099505 |
| FANCM | 1.39929425 | 0.01388292 |
| FAP | 2.770235 | 0.00086562 |
| FAR2 | 1.69657175 | 1.60E-05 |
| FAS | 2.27123475 | 0.002944624 |
| FASLG | 1.1360055 | 0.015096471 |
| FBLIM1 | 1.129112 | 0.000578739 |
| FBN1 | 1.5140595 | 0.000200771 |
| FBXL21P | -1.35146625 | 0.012691137 |
| FBXO16 | -1.06410275 | 0.000616183 |
| FBXO32 | 4.398911 | 1.67E-06 |
| FBXO40 | 3.97531125 | 1.81E-09 |
| FBXO48 | 1.15064625 | 0.000920062 |
| FCAMR | 1.701766 | 0.006803476 |
| FCER1A | 3.29832275 | 0.000259625 |
| FCER1G | 2.57070975 | 3.09E-05 |
| FCGBP | -1.46251675 | 0.000494658 |
| FCGR2B | 3.0617725 | 0.003392465 |
| FCGR3B | 1.30673525 | 0.003801291 |
| FCHO1 | 1.8146095 | 0.000718975 |
| FCMR | 2.804101 | 0.005294074 |
| FCRL3 | 1.866992 | 0.006494444 |
| FCRL6 | 2.99180675 | 0.004896376 |
| FERMT3 | 2.88269225 | 0.000367821 |
| FFAR2 | 3.20553575 | 0.00639414 |
| FGA | 4.64572 | 0.013884325 |
| FGD2 | 3.5821005 | 0.000115262 |
| FGF7 | 1.96956975 | 0.018770809 |
| FGF9 | -1.129628 | 0.000208988 |
| FGFBP1 | 2.539968 | 0.034027843 |
| FGL2 | 1.27243775 | 0.001334773 |
| FIBIN | 1.49532275 | 0.001507001 |
| FITM2 | -1.02114225 | 0.039059616 |
| FKBP5 | 3.08041075 | 7.03E-06 |
| FLI1 | 1.06476575 | 0.000511685 |
| FLT3 | 3.096635 | 0.00010952 |
| FLVCR2 | -1.19131725 | 0.000130207 |
| FMNL1 | 1.92961825 | 4.43E-05 |
| FN1 | 1.5426985 | 0.000424834 |
| FNDC4 | 3.32354725 | 0.00503151 |
| FOLR2 | -2.4285615 | 0.000158603 |
| FOS | 1.83042275 | 0.046805403 |
| FOSL1 | 5.73640725 | 0.000660598 |
| FOSL2 | 2.76258325 | 0.000114889 |
| FOXA3 | 3.36728325 | 0.011704593 |
| FOXM1 | 2.8265225 | 0.026443606 |
| FOXN1 | 1.814347 | 0.03137725 |
| FOXO1 | 1.114158 | 0.000426898 |
| FOXP3 | 2.354965 | 0.014979373 |
| FPR1 | 3.6967715 | 0.00126453 |
| FPR2 | 2.0746645 | 0.028980379 |
| FRMD6 | 1.104458 | 6.52E-05 |
| FRMD7 | -1.5760775 | 0.000494488 |
| FRMPD3 | 1.24157675 | 0.001464287 |
| FRRS1L | 2.439539 | 0.017084768 |
| FSCN1 | 1.67192875 | 0.001292355 |
| FUT1 | 1.544019 | 0.017368577 |
| FUT5 | 1.78906875 | 0.011338261 |
| FXYD5 | 2.156202 | 1.46E-05 |
| FXYD6 | 1.2725435 | 0.003007839 |
| FYN | 1.41111575 | 0.00062384 |
| FZD2 | -1.2983625 | 0.00072263 |
| FZD4 | 1.35023025 | 7.98E-06 |
| FZD8 | 1.356327 | 0.000221856 |
| G0S2 | -1.42921 | 0.000948463 |
| G6PC | 1.875474 | 0.002126873 |
| G6PC2 | -2.1785615 | 0.025439141 |
| G6PD | 1.04070875 | 0.004838292 |
| GAB2 | 1.201754 | 0.000715752 |
| GABRE | -1.60824 | 0.04205164 |
| GABRR1 | 1.40710625 | 0.007878985 |
| GAD2 | -1.7996055 | 0.008381708 |
| GADD45A | 2.043019 | 1.31E-05 |
| GADD45B | 1.554156 | 0.015567232 |
| GADD45G | 1.986153 | 3.55E-05 |
| GAL3ST1 | -1.6338545 | 0.010407245 |
| GAL3ST3 | -2.070926 | 0.002127447 |
| GALNT15 | 1.5203475 | 0.000203427 |
| GALNT16 | 1.610019 | 0.004875699 |
| GALNT5 | -1.5078035 | 0.025803323 |
| GALNTL5 | 3.410964 | 0.015410825 |
| GAREM1 | 1.02765525 | 0.001693903 |
| GAS2L3 | 3.940453 | 1.88E-05 |
| GAS7 | 1.095203 | 0.005679763 |
| GASK1A | 1.34744275 | 0.007637545 |
| GATA2 | -1.0268895 | 0.048347409 |
| GATA5 | -1.79292825 | 0.014913869 |
| GATA6 | 1.42915975 | 0.000180905 |
| GCG | 7.23963525 | 0.001511572 |
| GCKR | 3.9100615 | 8.57E-06 |
| GCLC | 1.865973 | 1.33E-06 |
| GCNT4 | 1.0774015 | 0.010987089 |
| GDF15 | 4.36855475 | 0.002912068 |
| GDF6 | 3.9053375 | 0.001727378 |
| GEN1 | 2.23797075 | 0.001219615 |
| GFI1 | 2.74681575 | 0.00691148 |
| GFPT2 | 2.34888125 | 0.001734185 |
| GFRA1 | 2.14517525 | 0.008266085 |
| GFRA2 | -1.55342725 | 7.18E-05 |
| GGN | 1.27228125 | 0.002711757 |
| GIMAP2 | 1.39123225 | 0.000360239 |
| GIMAP6 | 1.3014745 | 0.000554292 |
| GINS3 | 1.1338605 | 0.000240094 |
| GINS4 | 2.459279 | 0.000132042 |
| GJA3 | -1.38723775 | 0.018617584 |
| GJA9 | -2.39717875 | 0.005197507 |
| GJB6 | -2.85122475 | 5.15E-06 |
| GJC1 | 1.1872425 | 0.001591544 |
| GJD2 | -4.2065625 | 0.001025189 |
| GK5 | 1.104378 | 0.005189068 |
| GLDN | 7.4053365 | 0.000355883 |
| GLI3 | 1.29506025 | 0.000769292 |
| GLIS2 | -1.19403475 | 0.000505536 |
| GLRA3 | -2.76004425 | 0.011337117 |
| GLTPD2 | -1.115256 | 0.000237356 |
| GLUL | 2.01462325 | 0.000380054 |
| GLYATL2 | -1.7129545 | 0.005973051 |
| GLYCTK | -1.06446275 | 0.000378975 |
| GM2A | 3.9138305 | 9.90E-06 |
| GMFG | 1.6286525 | 0.000485008 |
| GMIP | 1.95619925 | 0.00338451 |
| GNA14 | 1.084364 | 0.020409033 |
| GNA15 | 2.27776325 | 0.000210316 |
| GNG2 | 1.39089975 | 0.001455898 |
| GNLY | 2.434458 | 0.036349364 |
| GP1BA | 1.50966475 | 0.010350072 |
| GP6 | 4.395767 | 7.16E-05 |
| GPC2 | 2.36794725 | 0.000959366 |
| GPLD1 | -1.94101075 | 0.000892282 |
| GPNMB | 10.09477375 | 2.98E-08 |
| GPR1 | 1.1286755 | 0.010056558 |
| GPR132 | 3.58797275 | 0.002574507 |
| GPR143 | 1.167229 | 0.000513698 |
| GPR152 | 2.0424815 | 0.012336762 |
| GPR160 | 1.07790525 | 0.001141782 |
| GPR171 | 2.81910375 | 8.63E-05 |
| GPR174 | 3.728215 | 0.002787523 |
| GPR183 | 3.607274 | 0.005377621 |
| GPR19 | 2.6860935 | 0.022798522 |
| GPR3 | 1.308515 | 0.001893988 |
| GPR34 | 1.50493875 | 0.000723599 |
| GPR65 | 2.66373075 | 0.000177958 |
| GPR68 | 2.36999525 | 0.011652889 |
| GPR75 | 1.073952 | 0.049238919 |
| GPR82 | 2.9214575 | 0.044626882 |
| GPR83 | 1.95308425 | 0.009496747 |
| GPR85 | -1.99432 | 0.046666035 |
| GPRC5A | 1.80801275 | 0.002171784 |
| GPRC6A | -1.94761 | 0.00016509 |
| GRAP | 3.6040945 | 0.000252664 |
| GRAP2 | 1.381092 | 0.03203984 |
| GREB1 | 1.26399 | 0.004875927 |
| GRIA1 | -2.89624075 | 0.019177151 |
| GRID2 | 1.414451 | 0.041344398 |
| GRIN2B | -1.261619 | 0.000441426 |
| GRK3 | 2.23057175 | 5.61E-06 |
| GRM2 | 1.55036675 | 0.013428466 |
| GRM5 | -1.19426825 | 0.027704465 |
| GRPR | 2.241446 | 0.014421742 |
| GTSE1 | 3.06715725 | 0.013488464 |
| GUCA2A | -1.49472175 | 0.012693269 |
| GUCA2B | -1.5501565 | 0.017786198 |
| GYPC | 1.4933365 | 0.000124215 |
| GZMA | 3.3382015 | 0.009250602 |
| GZMH | 1.672609 | 0.014749216 |
| GZMK | 3.967301 | 3.89E-06 |
| HABP2 | 4.06567075 | 8.04E-05 |
| HAPLN3 | 1.53104075 | 0.011696731 |
| HASPIN | 3.40498075 | 0.013260658 |
| HAVCR1 | 2.88742025 | 0.003894492 |
| HAVCR2 | 1.62061525 | 0.000159619 |
| HBEGF | 1.8267865 | 0.010378206 |
| HCAR1 | -1.51836325 | 0.034413567 |
| HCLS1 | 2.52517525 | 3.46E-05 |
| HCST | 2.7566105 | 4.84E-05 |
| HDAC4 | 1.02907375 | 0.00101776 |
| HDC | -3.44512825 | 0.000465572 |
| HEMGN | 5.2779075 | 0.011974511 |
| HEMK1 | -1.134772 | 0.000463116 |
| HENMT1 | -2.56968775 | 4.00E-06 |
| HERPUD1 | 2.35998925 | 1.10E-06 |
| HES2 | 1.42272475 | 0.011897554 |
| HESX1 | -1.6193385 | 0.001337549 |
| HEXB | 1.28964975 | 0.000243792 |
| HEY1 | 1.445392 | 0.000380339 |
| HGF | 1.13820825 | 0.003524346 |
| HHEX | 1.0490605 | 0.008510734 |
| HHIPL1 | 2.0943905 | 0.007090687 |
| HIF3A | 2.1595875 | 0.001496326 |
| HIGD1B | 1.16393925 | 0.011196289 |
| HIVEP2 | 1.2773885 | 0.000573173 |
| HJURP | 1.11222525 | 0.001197257 |
| HK2 | 4.7831795 | 0.000614785 |
| HK3 | 3.8645895 | 0.000221415 |
| HMCN2 | 2.5307395 | 0.002527303 |
| HMGA1 | 1.7411385 | 0.005127807 |
| HMGB2 | 1.347628 | 0.003149116 |
| HMGB3 | 1.6696605 | 0.004608007 |
| HMGCS2 | -1.40664225 | 0.000970214 |
| HMMR | 1.19707125 | 0.043323241 |
| HMOX1 | 4.64173825 | 1.06E-06 |
| HMX2 | -2.896331 | 0.006268498 |
| HNF1A | -1.13349975 | 0.000301735 |
| HORMAD1 | 3.23063025 | 0.001335083 |
| HOXB6 | -1.17445025 | 0.000531592 |
| HOXC5 | -1.13813 | 0.002283142 |
| HOXC6 | -1.069982 | 0.012943255 |
| HOXC9 | -1.45872975 | 4.65E-05 |
| HOXD10 | -1.1211505 | 0.005633933 |
| HOXD3 | -1.224963 | 0.000636091 |
| HPCAL1 | 1.22169675 | 0.000270692 |
| HRH1 | 1.35254375 | 0.004204731 |
| HRH2 | 2.2524425 | 0.000782872 |
| HS3ST1 | 1.89637675 | 0.003671612 |
| HSPB1 | 1.35662275 | 0.005505001 |
| HSPB7 | 1.3200245 | 0.014456856 |
| HSPB8 | 1.342301 | 0.002604847 |
| HTR1F | -1.3851775 | 0.039276957 |
| HTR2A | -2.85788875 | 2.02E-05 |
| HTRA3 | 1.6929105 | 5.12E-05 |
| HVCN1 | 1.173932 | 0.00814086 |
| HYPM | 2.3349625 | 0.011629673 |
| IAPP | -2.241446 | 0.014421742 |
| ICAM1 | 2.08936025 | 0.001011982 |
| ICAM3 | 2.11532675 | 9.21E-06 |
| ICAM5 | 4.51373575 | 0.000848159 |
| ICOS | 4.15374975 | 9.96E-05 |
| IDO1 | 5.6212385 | 4.28E-05 |
| IER5 | 1.05213675 | 0.001059899 |
| IFNLR1 | 1.0918465 | 0.019963135 |
| IFRD1 | 1.5919895 | 0.002599396 |
| IGFBP1 | 4.78171875 | 0.001155292 |
| IGFBP4 | 1.046438 | 0.001619193 |
| IGSF23 | -1.24284625 | 0.005322667 |
| IGSF6 | 1.54724625 | 0.002174997 |
| IGSF9 | 3.22750475 | 0.027231352 |
| IKBKE | 1.6382605 | 0.00071492 |
| IKZF1 | 1.377077 | 0.002116373 |
| IKZF3 | 4.528781 | 0.002297605 |
| IL10RA | 2.923947 | 1.13E-05 |
| IL11 | 3.781333 | 0.005917156 |
| IL12A | 1.824802 | 0.046944019 |
| IL12B | -1.06019275 | 0.006452714 |
| IL13RA1 | 1.0772645 | 7.91E-05 |
| IL15RA | 1.23133925 | 0.00392215 |
| IL17RB | 2.46649825 | 1.19E-06 |
| IL18BP | 4.0456345 | 2.15E-07 |
| IL18RAP | 1.39305875 | 0.016733947 |
| IL1R2 | 5.21532275 | 0.001057241 |
| IL1RL1 | 7.3206975 | 6.22E-06 |
| IL24 | 2.01472325 | 0.002477117 |
| IL27RA | 1.228431 | 0.000729373 |
| IL2RA | 2.4651505 | 0.008143754 |
| IL2RB | 1.664235 | 0.000184667 |
| IL2RG | 5.04782425 | 7.38E-05 |
| IL4I1 | 2.16692325 | 0.000212296 |
| IL4R | 2.72194 | 3.57E-05 |
| IL6 | 3.00448075 | 0.012506215 |
| IL7R | 5.83192825 | 5.89E-05 |
| IL9 | -1.70618275 | 0.031134585 |
| INAVA | 1.53071875 | 0.042082541 |
| INHBB | 3.7701725 | 1.23E-05 |
| INKA1 | -1.87200775 | 0.022782734 |
| INPP5D | 2.80901325 | 1.03E-05 |
| INSYN2 | -1.68428325 | 1.39E-05 |
| IP6K2 | 1.3213845 | 0.001525762 |
| IQCA1 | 3.0598665 | 0.003577267 |
| IQCM | -1.500531 | 0.001250394 |
| IQCN | 1.44426375 | 0.017076038 |
| IQGAP3 | 1.25987675 | 0.003404884 |
| IRF1 | 1.11129975 | 0.023233102 |
| IRF4 | 3.15752825 | 0.033863788 |
| IRF5 | 1.54085475 | 0.006579751 |
| IRF8 | 2.160501 | 7.22E-05 |
| IRS1 | 1.6394515 | 1.59E-06 |
| IRS2 | 2.35419325 | 3.36E-05 |
| IRX5 | -1.4027135 | 2.82E-05 |
| ISG20 | 1.64269925 | 0.002905331 |
| ISLR2 | 2.28514325 | 0.003716996 |
| ITGA4 | 2.34003325 | 0.000128758 |
| ITGA5 | 2.22980475 | 1.15E-05 |
| ITGAD | 2.975215 | 2.82E-06 |
| ITGAM | 4.23161675 | 1.01E-06 |
| ITGAX | 6.86522075 | 1.09E-06 |
| ITGB7 | 1.92175925 | 1.54E-05 |
| ITGB8 | 1.16656375 | 0.007429807 |
| ITIH1 | 2.24432025 | 0.000319604 |
| ITIH4 | 1.0110895 | 0.000345245 |
| ITK | 3.14771625 | 1.60E-06 |
| ITPKC | 1.857659 | 1.47E-06 |
| ITPRIP | 1.13749275 | 0.002268261 |
| JAG1 | -1.2839305 | 5.27E-05 |
| JAK3 | 4.378483 | 1.63E-05 |
| JAKMIP1 | 1.88722825 | 0.000671783 |
| JAML | 2.50600175 | 2.71E-05 |
| JCHAIN | 4.16031775 | 0.000526542 |
| JHY | 3.671406 | 0.002875645 |
| JUN | 1.70725275 | 0.014141464 |
| JUNB | 2.70560675 | 0.003197689 |
| KANK3 | 1.13073775 | 0.002214003 |
| KANK4 | -1.58235925 | 8.32E-05 |
| KBTBD7 | -1.8412175 | 0.011969432 |
| KCNA1 | 1.6823125 | 0.013926705 |
| KCNA3 | -1.10803875 | 0.004763219 |
| KCNA4 | -1.93872225 | 0.011873962 |
| KCNA6 | 2.688722 | 0.005730169 |
| KCNAB2 | 2.74533625 | 7.17E-06 |
| KCND1 | 1.444107 | 0.000868915 |
| KCNE2 | -1.03953975 | 0.000834804 |
| KCNE3 | 4.118194 | 7.85E-05 |
| KCNJ13 | -1.4322025 | 0.000102553 |
| KCNJ2 | 1.53107325 | 0.000417945 |
| KCNJ9 | -1.08915075 | 0.009199963 |
| KCNK2 | -1.65558475 | 6.63E-05 |
| KCNK6 | 1.196777 | 0.00483569 |
| KCNMA1 | -2.04116675 | 0.000173797 |
| KCNMB4 | -1.21729325 | 0.00385455 |
| KCTD14 | -1.0199815 | 0.016272342 |
| KCTD15 | -1.61333375 | 0.000276609 |
| KCTD21 | -1.3056915 | 0.000225529 |
| KCTD7 | -1.86957225 | 0.000140892 |
| KDR | -1.43152825 | 0.000568419 |
| KERA | 2.18994325 | 0.035342716 |
| KIAA1549 | 1.33628325 | 0.000683955 |
| KIAA1755 | 3.272746 | 1.47E-06 |
| KIF11 | 3.03135925 | 0.003468249 |
| KIF14 | 3.22665825 | 0.008677903 |
| KIF15 | 2.40012125 | 0.000236586 |
| KIF18B | 3.00236975 | 0.038997306 |
| KIF1A | 2.34506125 | 0.011818263 |
| KIF20A | 3.645099 | 0.015238085 |
| KIF21B | 1.999973 | 0.001705027 |
| KIF22 | 1.41050725 | 0.006232235 |
| KIF23 | 1.59052375 | 0.002196529 |
| KIF26A | -1.2349185 | 0.000595823 |
| KIF2C | 3.089382 | 0.01561833 |
| KIF3C | 1.29828375 | 0.007138241 |
| KIFC1 | 1.02090575 | 0.006068684 |
| KISS1 | -1.0664735 | 0.008907698 |
| KL | -1.09879825 | 0.000707913 |
| KLF10 | 1.6951815 | 0.000392144 |
| KLF11 | 1.3529765 | 0.000188334 |
| KLF13 | 2.48675575 | 0.02196177 |
| KLF15 | 1.128382 | 0.016501119 |
| KLF2 | -1.19498625 | 0.000664488 |
| KLF4 | 1.1915315 | 0.004422062 |
| KLF6 | 1.731927 | 0.000418088 |
| KLF7 | 1.02310725 | 0.000868393 |
| KLF9 | 1.2622805 | 0.000132845 |
| KLHDC7A | -1.09120975 | 0.00037882 |
| KLHL25 | 1.08135075 | 0.000120258 |
| KLHL29 | 1.35745625 | 0.001708505 |
| KLHL3 | -1.06657225 | 0.028505751 |
| KLHL6 | 2.1956455 | 0.000356739 |
| KLK1 | -1.07486425 | 0.022018434 |
| KLRF1 | 1.58220725 | 0.018333028 |
| KLRG2 | 1.24907725 | 0.005026196 |
| KNL1 | 1.5169525 | 0.028571451 |
| KNTC1 | 1.2697775 | 0.012343033 |
| KPNA5 | -1.49430525 | 0.000156639 |
| KRT7 | 1.24571375 | 0.028204687 |
| KYAT1 | 1.150622 | 0.000520098 |
| LACRT | -1.2425155 | 0.01267163 |
| LAIR1 | 5.3625565 | 3.48E-05 |
| LAMB4 | -1.3594305 | 0.017600761 |
| LAMC2 | 1.26410875 | 0.001901318 |
| LAMP3 | 2.0978925 | 0.034673767 |
| LANCL3 | 1.427821 | 0.001201603 |
| LAPTM5 | 3.05854325 | 4.71E-05 |
| LAT2 | 1.4116475 | 0.003481968 |
| LAX1 | 1.58241375 | 0.005960358 |
| LAYN | 1.1819305 | 0.017912673 |
| LBHD2 | 2.63648225 | 0.018392023 |
| LCAT | 1.26486225 | 0.013922273 |
| LCK | 1.16227275 | 0.005724705 |
| LCN2 | 2.34072225 | 0.00738377 |
| LCP1 | 1.195444 | 0.002451616 |
| LCP2 | 4.097497 | 2.74E-06 |
| LECT2 | -1.234062 | 0.005027788 |
| LEP | -2.69010325 | 0.009104878 |
| LGALS3 | 2.35834075 | 0.000118419 |
| LGR6 | -2.74748675 | 0.025313394 |
| LHFPL1 | -2.85824 | 0.004990706 |
| LHX1 | -1.65689775 | 0.000352179 |
| LIF | 3.41065525 | 0.000793824 |
| LIMD2 | 1.7229835 | 0.000216799 |
| LIMK1 | 1.258193 | 0.000525542 |
| LIPJ | 2.60322675 | 0.043346712 |
| LMLN2 | -1.2482165 | 0.022141395 |
| LMX1A | -3.51323375 | 0.001364121 |
| LNP1 | -2.1498755 | 0.020177786 |
| LONRF3 | 1.616118 | 1.35E-05 |
| LOX | 1.243205 | 0.022658393 |
| LOXL2 | 1.5895665 | 0.000126054 |
| LPAR2 | 1.80812775 | 0.01455948 |
| LPAR3 | 1.4615615 | 0.001920579 |
| LPAR5 | 2.2379945 | 0.01565736 |
| LPCAT1 | -1.8556425 | 1.83E-05 |
| LPCAT2 | 1.77959175 | 0.000198856 |
| LPXN | 1.88703475 | 5.28E-05 |
| LRG1 | 1.762524 | 0.014192205 |
| LRIG3 | -1.3777235 | 0.001244084 |
| LRMP | 1.9425545 | 0.001920385 |
| LRP1 | 1.4416005 | 0.000475664 |
| LRP8 | 2.49932775 | 6.81E-06 |
| LRRC10B | 1.20553725 | 0.012425468 |
| LRRC17 | -3.05445725 | 4.84E-05 |
| LRRC23 | 2.26344375 | 0.002506004 |
| LRRC38 | 1.77365275 | 0.009296059 |
| LRRC3B | -2.599317 | 0.000155438 |
| LRRC66 | -1.1918055 | 0.009544332 |
| LRRC74B | 2.51920425 | 8.76E-08 |
| LRRIQ1 | 1.68526025 | 0.002299013 |
| LRRK2 | -1.39016975 | 0.000120708 |
| LRRN1 | -1.08488775 | 0.021575861 |
| LRRN3 | -3.72502825 | 2.30E-05 |
| LRRTM1 | -1.27196825 | 0.02437963 |
| LRRTM3 | -3.007753 | 0.022269262 |
| LSP1 | 1.187466 | 0.000669593 |
| LST1 | 2.3597435 | 0.005188309 |
| LTB4R | 1.207129 | 0.012174976 |
| LTB4R2 | 1.97672275 | 0.016757067 |
| LTBP2 | 2.395702 | 0.001392961 |
| LTF | 2.23125225 | 0.000314048 |
| LUM | 2.75654375 | 0.000512579 |
| LVRN | -3.93809525 | 0.000199538 |
| LY6E | 1.01139975 | 0.002411811 |
| LYG2 | 1.9534455 | 0.003443571 |
| LYPD6 | -1.70947875 | 0.000567948 |
| LYVE1 | 1.2828585 | 0.00272921 |
| M1AP | -1.70067625 | 0.000365029 |
| MAB21L3 | 2.47462275 | 2.77E-05 |
| MAB21L4 | 2.06736525 | 0.018581142 |
| MADCAM1 | 2.5147605 | 0.010174351 |
| MAFF | 4.52158525 | 2.42E-05 |
| MAG | 2.269204 | 0.012809891 |
| MAK | 2.21651625 | 0.000469696 |
| MAN1C1 | 1.14430725 | 0.001471827 |
| MANSC4 | -1.08444275 | 0.007809893 |
| MAP2K6 | -1.002301 | 0.002460046 |
| MAP3K14 | 1.0887185 | 0.00025771 |
| MAP3K19 | 2.30720475 | 0.012978852 |
| MAP3K8 | 2.208578 | 0.000710999 |
| MAP3K9 | -1.03534625 | 0.000611311 |
| MAP4K1 | 2.6326185 | 0.001707679 |
| MAPK6 | 1.6767345 | 8.92E-06 |
| MASP1 | 3.49888275 | 0.003055948 |
| MASTL | 3.02997125 | 0.018627632 |
| MATK | 1.6815255 | 0.039742521 |
| MATN4 | -1.19713275 | 0.01278682 |
| MBLAC2 | -1.0415235 | 0.001209738 |
| MCEMP1 | 4.3598605 | 3.43E-07 |
| MCM5 | 1.45264775 | 0.000382983 |
| MDM2 | 1.43004925 | 1.98E-05 |
| MEDAG | 2.131738 | 0.013068261 |
| MEF2C | 1.14851775 | 0.000218498 |
| MEFV | 4.28250175 | 0.000186677 |
| MEI4 | 2.71367825 | 0.021072077 |
| MEIS2 | -1.21238825 | 0.000138373 |
| MELK | 1.91441125 | 0.049873831 |
| MEOX1 | 1.881425 | 0.005267141 |
| MESP2 | -1.949994 | 0.016755067 |
| MEX3A | 2.5859005 | 0.009838257 |
| MEX3B | -1.36416525 | 0.008275981 |
| MFRP | 3.1469445 | 0.000523564 |
| MFSD2A | 2.99007725 | 2.70E-05 |
| MFSD9 | -1.23291575 | 0.041503609 |
| MGARP | -1.86344325 | 0.034178315 |
| MGAT2 | -1.10329175 | 0.000685302 |
| MGAT3 | -1.4073795 | 0.000281223 |
| MICAL1 | 1.475534 | 5.19E-05 |
| MILR1 | 2.1662635 | 7.31E-05 |
| MIR103A-2 | -1.2924815 | 0.014674946 |
| MIR142 | 1.6536775 | 0.017131903 |
| MIR147B | 2.06997175 | 0.000208069 |
| MIR15A | -2.16808825 | 0.044991992 |
| MIR191 | 1.60375925 | 0.002935156 |
| MIR200A | -1.817309 | 0.000208475 |
| MIR21 | 1.789634 | 0.000863276 |
| MIR22 | 2.2268935 | 0.002512769 |
| MIR34A-1 | 1.051414 | 0.012456972 |
| MIR365B | -2.42070775 | 0.024415552 |
| MIR450A-1 | -2.3592065 | 0.000255408 |
| MIR450A-2 | -1.98518875 | 0.012762372 |
| MIR6529 | -1.18872225 | 0.013620296 |
| MIR9771F | -1.2924815 | 0.049849561 |
| MIR9771K | 1.14624075 | 0.023107597 |
| MIS18BP1 | 2.27545075 | 0.004846011 |
| MITD1 | -1.27496725 | 0.002737352 |
| MKI67 | 2.7822595 | 0.001882252 |
| MKRN2OS | -1.2138305 | 0.000236872 |
| MKRN3 | -2.08496275 | 0.012819002 |
| MLNR | 3.39056075 | 1.42E-05 |
| MMD | 2.7817165 | 0.000211994 |
| MMP1 | 9.0677855 | 3.30E-05 |
| MMP12 | 8.355371 | 7.85E-06 |
| MMP13 | 3.47399875 | 0.004294184 |
| MMP14 | 2.0108395 | 7.79E-05 |
| MMP19 | 1.75207975 | 1.51E-05 |
| MMP2 | 1.3553605 | 4.22E-05 |
| MMP23B | 1.398129 | 0.012757016 |
| MMP25 | 1.66943 | 0.004306049 |
| MMP7 | 4.42180375 | 3.24E-06 |
| MMP8 | 3.074802 | 0.011561309 |
| MMP9 | 5.717146 | 0.000100612 |
| MMRN2 | 1.010989 | 0.005306768 |
| MNT | 1.201347 | 0.000102787 |
| MOBP | 2.9823145 | 0.004590589 |
| MOG | 2.392856 | 6.62E-06 |
| MORN3 | 2.31175575 | 0.010556377 |
| MPEG1 | 1.9909935 | 0.000154341 |
| MPP2 | -1.34386975 | 0.034925406 |
| MPPED1 | 2.65699025 | 0.003477634 |
| MRC1 | 2.52667775 | 1.24E-05 |
| MS4A10 | -1.02126575 | 0.004162352 |
| MS4A14 | 2.519204 | 0.006566568 |
| MS4A3 | 4.2572195 | 5.45E-06 |
| MS4A4A | 2.24842325 | 8.57E-05 |
| MSR1 | 3.34054775 | 9.73E-08 |
| MST1 | 1.70497825 | 0.045035057 |
| MST1R | 3.4298365 | 0.000961257 |
| MTBP | 2.62423775 | 0.000748475 |
| MTHFD2 | 1.678106 | 0.000108896 |
| MTNR1A | 1.25236725 | 0.001418803 |
| MUC13 | 4.65171775 | 0.000336383 |
| MUC6 | 1.5118715 | 0.014260635 |
| MXRA5 | 3.92225125 | 0.00015279 |
| MYBL2 | 4.05074925 | 0.006238874 |
| MYBPC1 | 1.65422075 | 0.012616797 |
| MYBPC2 | 1.31120275 | 0.000155069 |
| MYC | 3.2625815 | 8.13E-05 |
| MYCN | -1.622614 | 3.84E-05 |
| MYL2 | 4.738262 | 1.60E-08 |
| MYO1F | 2.1772975 | 0.000151664 |
| MYO1G | 1.9249455 | 0.011505508 |
| MZB1 | 1.88396675 | 0.003432484 |
| NABP1 | 1.98579475 | 1.45E-05 |
| NAP1L3 | -1.481238 | 0.000173335 |
| NAPSA | 1.23579625 | 0.00237512 |
| NAT16 | 4.509349 | 0.001752635 |
| NAV3 | 1.035215 | 0.005249012 |
| NCAPH | 4.64714575 | 9.88E-07 |
| NCF1 | 3.40266325 | 0.000227383 |
| NCF2 | 3.522069 | 0.000202666 |
| NCF4 | 1.6805475 | 0.006198083 |
| NCKAP1L | 2.5198215 | 0.000200815 |
| NCOA7 | -1.55852275 | 0.00698163 |
| NCR3LG1 | 2.39171125 | 7.59E-06 |
| NDEL1 | 1.02421275 | 0.001597527 |
| NECAB2 | 3.4538485 | 0.00045629 |
| NECAB3 | -2.6033045 | 0.009697428 |
| NEK2 | 1.68925325 | 0.017723077 |
| NET1 | 1.22112675 | 0.00101414 |
| NETO2 | 2.25349875 | 0.010845537 |
| NEURL1B | -2.620507 | 0.030711824 |
| NFAM1 | 1.311271 | 0.036636602 |
| NFE2L3 | 1.05292975 | 0.01139837 |
| NFIL3 | 1.51561325 | 0.010985854 |
| NFKBIA | 1.7460735 | 0.005889947 |
| NFKBID | 1.91194125 | 0.001081288 |
| NFKBIZ | 1.77550725 | 0.002498762 |
| NKG7 | 2.856129 | 0.008852518 |
| NLGN3 | -2.57480175 | 0.006895112 |
| NLRC3 | -2.057962 | 0.00015207 |
| NLRC5 | 1.016588 | 0.000411614 |
| NLRP12 | 2.519204 | 0.015718138 |
| NLRP3 | 3.2054815 | 1.24E-05 |
| NLRP6 | -2.20352525 | 2.08E-05 |
| NMNAT3 | -1.55866075 | 0.000689278 |
| NMUR1 | 2.82700325 | 0.007870012 |
| NNAT | 1.193899 | 0.017723522 |
| NNMT | 2.877444 | 0.006351962 |
| NOA1 | -1.3594145 | 2.70E-05 |
| NOS1 | -1.16404725 | 0.007064399 |
| NOSTRIN | 1.39062075 | 6.25E-05 |
| NPAS2 | 1.7411345 | 5.03E-06 |
| NPAS4 | 2.78456775 | 0.004365839 |
| NPHS2 | -1.28578875 | 0.004250751 |
| NPTX1 | 2.466562 | 0.005927975 |
| NPTX2 | 2.241446 | 0.014421742 |
| NPY4R2 | 1.58048175 | 0.001762407 |
| NR0B1 | 1.8415805 | 0.04646979 |
| NR0B2 | -1.41612275 | 0.000718681 |
| NR1D1 | 1.16742225 | 6.36E-05 |
| NR1H4 | -1.55097825 | 0.000248029 |
| NR2E1 | -1.03932425 | 0.023322745 |
| NR4A1 | 2.0702525 | 0.013581851 |
| NR4A3 | 3.07402025 | 0.011655472 |
| NREP | -1.0533905 | 0.000227942 |
| NRSN1 | -1.018577 | 0.000572105 |
| NT5C1B | -1.71375575 | 0.036023791 |
| NT5C3A | 1.0706995 | 0.007503605 |
| NT5DC4 | 5.32807 | 6.71E-07 |
| NTF3 | -1.6935835 | 0.000133588 |
| NTM | 1.39217175 | 0.003088211 |
| NUAK2 | -1.41365175 | 8.56E-05 |
| NUDT4 | -1.119635 | 0.000264112 |
| NUF2 | 2.50248625 | 0.003011322 |
| NUP62 | 1.19784175 | 0.002735411 |
| NUSAP1 | 1.798824 | 0.010423497 |
| NXPE3 | 2.814058 | 0.000858268 |
| NXPH2 | -2.097495 | 0.00765421 |
| NXPH3 | 1.01680675 | 0.008150319 |
| NXPH4 | -1.5842335 | 0.006241036 |
| NYAP2 | 4.084336 | 3.63E-05 |
| OBSL1 | -1.00433925 | 0.007427006 |
| OIP5 | 2.73298 | 0.012548733 |
| OIT3 | -1.97475275 | 5.80E-05 |
| OLFM4 | 2.9257935 | 0.000220955 |
| OLFML2A | 1.02180325 | 0.004073909 |
| OLFML3 | 1.126807 | 0.004193667 |
| OLIG2 | 1.8729635 | 0.01221658 |
| OMG | 3.39617375 | 0.000325566 |
| OPCML | 1.79248125 | 0.024440433 |
| OPTC | 2.80358 | 0.005584837 |
| OSCAR | 2.93809525 | 0.00216001 |
| OSM | 3.96845 | 0.000537422 |
| OSR1 | -4.7400345 | 1.53E-07 |
| OSR2 | -2.031073 | 0.007320301 |
| OVCH2 | -2.6643165 | 0.034567528 |
| OVOL1 | -1.12369425 | 0.000128658 |
| OXGR1 | -1.44303 | 0.00125604 |
| P2RY13 | 1.4099855 | 0.035497604 |
| P2RY14 | 2.306457 | 1.32E-05 |
| P4HA3 | 2.269204 | 0.012809891 |
| PACRG | 1.42481075 | 0.010898372 |
| PADI4 | 3.4518115 | 0.00949877 |
| PAK3 | -1.03561825 | 0.015211547 |
| PALB2 | 1.81899725 | 0.000102112 |
| PALD1 | 1.3805755 | 0.00015703 |
| PANX1 | 1.026661 | 0.011263554 |
| PARM1 | 1.003456 | 0.019377038 |
| PARPBP | 2.59006625 | 0.020818036 |
| PARVG | 3.96681575 | 3.84E-05 |
| PBK | 3.6114265 | 0.016977621 |
| PBX4 | 1.92856125 | 0.038984988 |
| PCDH12 | -1.77711475 | 0.005605168 |
| PCDH18 | -1.24277475 | 0.001253673 |
| PCED1B | -1.57858975 | 0.000297587 |
| PCK1 | 1.10288125 | 0.002070172 |
| PCLAF | 1.60749025 | 0.000211247 |
| PCNX4 | -1.11138125 | 0.000490593 |
| PCSK1 | 5.57151825 | 0.000213634 |
| PDCD1 | 3.5560005 | 0.004549677 |
| PDCD1LG2 | 4.1683605 | 0.00021276 |
| PDE10A | 2.2681765 | 0.000829503 |
| PDE2A | 1.0424065 | 0.000309123 |
| PDE4B | 2.516176 | 6.25E-05 |
| PDE4C | 1.40635575 | 0.00262162 |
| PDE6A | 1.538597 | 0.015572866 |
| PDE6H | 1.12687225 | 0.041090651 |
| PDK4 | 3.576537 | 0.000100503 |
| PDLIM1 | 2.09212325 | 6.95E-05 |
| PDLIM4 | 2.2070855 | 0.012447151 |
| PDLIM7 | 1.12646775 | 0.000136621 |
| PDP1 | 1.33242325 | 0.000888078 |
| PDPN | 2.96230125 | 0.000410452 |
| PDYN | 4.015367 | 0.00330411 |
| PDZD2 | 1.22747025 | 0.006136029 |
| PEA15 | 1.02889125 | 0.031012661 |
| PER1 | 1.90683525 | 7.94E-05 |
| PER3 | -1.097541 | 0.009882907 |
| PET117 | -1.2429265 | 0.030862061 |
| PEX11A | -1.30884875 | 0.010368897 |
| PFKFB3 | 2.13686725 | 0.007500168 |
| PFN3 | -2.525822 | 0.018479076 |
| PGLYRP1 | 2.5510665 | 0.024342892 |
| PGP | -1.1675895 | 0.016101122 |
| PHLDA2 | 1.72171475 | 0.015396226 |
| PHTF2 | 1.6320225 | 0.001656267 |
| PHYHIPL | -2.39510375 | 0.003479202 |
| PI15 | 5.0137155 | 4.36E-05 |
| PIEZO1 | 1.060554 | 0.000495113 |
| PIF1 | 2.9285615 | 7.02E-05 |
| PIGL | -1.077647 | 0.006897373 |
| PIK3AP1 | 1.80891875 | 0.002755879 |
| PIK3CD | 1.17707475 | 0.006668972 |
| PIK3CG | 1.205683 | 0.000798754 |
| PIK3R2 | -1.02255475 | 0.001499011 |
| PIK3R3 | -1.08582 | 0.000230725 |
| PIK3R5 | 4.09571375 | 1.66E-06 |
| PILRA | 2.9821815 | 0.000659611 |
| PIM1 | 2.206253 | 0.000435593 |
| PIP4P2 | -1.09040475 | 0.005373909 |
| PIP5KL1 | 4.015668 | 0.000577898 |
| PITPNC1 | 1.87719225 | 0.000300185 |
| PKMYT1 | 1.179912 | 0.003311913 |
| PLA2G12B | -2.30416075 | 0.033895547 |
| PLA2G1B | -1.9036775 | 0.01445976 |
| PLA2G2A | 2.04341925 | 0.001238988 |
| PLA2G2D | 3.65588475 | 3.99E-06 |
| PLA2G4C | 1.0658205 | 0.000839784 |
| PLAT | 2.45712775 | 0.000131556 |
| PLAUR | 3.4340035 | 1.36E-05 |
| PLCB2 | 2.531086 | 2.01E-05 |
| PLEK | 4.11119 | 1.30E-05 |
| PLEKHF1 | 1.00487975 | 0.004441058 |
| PLEKHH2 | -1.11720925 | 0.000495256 |
| PLEKHO1 | 1.316374 | 0.000219619 |
| PLEKHO2 | 1.73142725 | 0.000427306 |
| PLIN1 | -1.210206 | 0.004057884 |
| PLIN5 | 1.53655875 | 0.001600654 |
| PLK3 | 1.7024705 | 0.002451029 |
| PLK4 | 1.25333675 | 0.000154321 |
| PLP2 | 1.11219675 | 0.000880823 |
| PLXNA2 | 1.19198775 | 0.000437291 |
| PLXNC1 | 2.263804 | 0.001032822 |
| PMAIP1 | 2.03743675 | 0.003818588 |
| PMCH | 1.976932 | 0.002331105 |
| PNMA2 | 3.56567075 | 0.002176261 |
| PNMT | 5.65711225 | 0.00041432 |
| PNPLA2 | 1.29567525 | 0.000351647 |
| PNRC1 | 1.46690825 | 5.00E-05 |
| POC1A | 1.05447725 | 0.002852841 |
| POLR1A | -1.463924 | 0.002384769 |
| POPDC2 | 1.23488025 | 0.014346974 |
| POSTN | 2.13512225 | 0.000269154 |
| POU2AF1 | 3.5397355 | 7.32E-06 |
| PPP1R15A | 1.52212325 | 0.005697361 |
| PPP1R1C | -1.1014645 | 0.004762661 |
| PPTC7 | 1.5022845 | 0.00017009 |
| PRAM1 | 4.12351675 | 0.010812609 |
| PRDM1 | 3.12965675 | 1.78E-06 |
| PRDM2 | 1.438707 | 8.64E-05 |
| PREX1 | 1.23072325 | 0.000804527 |
| PRF1 | 2.828582 | 0.000608488 |
| PRG2 | 2.9985885 | 0.019502076 |
| PRKAG3 | 1.18938275 | 0.002436762 |
| PRKCB | 1.8701845 | 0.008866019 |
| PRKCG | 1.213174 | 0.000796967 |
| PRLR | -1.398874 | 0.002206238 |
| PROX1 | -2.200917 | 0.000877793 |
| PRPH | 3.310481 | 0.002912413 |
| PRR15 | -1.297919 | 0.000766573 |
| PRR16 | 1.793439 | 0.004438703 |
| PRR35 | -1.8668965 | 0.000139891 |
| PRRX1 | 1.012223 | 0.01217569 |
| PRRX2 | 1.21763525 | 0.002963773 |
| PRSS12 | 3.40742075 | 4.00E-05 |
| PRSS36 | -1.14647525 | 0.047795437 |
| PRTFDC1 | 1.0387495 | 0.038009803 |
| PSD4 | 1.19272225 | 0.042833247 |
| PSMB9 | 1.28766225 | 0.000759241 |
| PSORS1C2 | 2.09424225 | 0.022394414 |
| PSTPIP1 | 2.0406065 | 0.000394978 |
| PTAFR | 1.680938 | 0.007663405 |
| PTGDR | -1.0156225 | 0.020134479 |
| PTGER2 | 1.93444025 | 5.69E-05 |
| PTGFR | -1.268774 | 0.035436709 |
| PTGIS | 1.28169825 | 0.031546103 |
| PTGR1 | 1.022057 | 0.023046737 |
| PTGS2 | 2.32090275 | 0.024361863 |
| PTH2R | -1.33614 | 0.006597486 |
| PTPDC1 | -1.68810925 | 0.005386501 |
| PTPN1 | 1.94229775 | 0.000238208 |
| PTPN7 | 2.50758025 | 0.00117096 |
| PTPRC | 2.6598375 | 2.39E-05 |
| PTPRE | 2.0068655 | 9.87E-05 |
| PTPRR | -2.72680225 | 9.63E-05 |
| PWWP3B | -1.2242125 | 0.001436697 |
| PXDNL | 2.891672 | 0.01795201 |
| PXT1 | 2.20183875 | 0.011748581 |
| PYCARD | 2.36488725 | 1.31E-05 |
| PYCR1 | 1.1632315 | 0.007066124 |
| PYDC1 | 2.97766075 | 4.66E-06 |
| QRICH2 | 5.5881065 | 0.00505517 |
| RAB38 | -1.11382575 | 0.017631076 |
| RAB39B | 1.47965875 | 0.006589585 |
| RAB3B | 2.05075475 | 0.027765011 |
| RAB40B | 2.01618575 | 0.041488614 |
| RAB7B | 5.086843 | 8.88E-06 |
| RAC2 | 3.13788225 | 2.62E-05 |
| RAD51 | 1.57217025 | 0.000442128 |
| RAD54L | 2.79406275 | 0.01366175 |
| RAG2 | -1.0926735 | 0.018882034 |
| RAI14 | 1.0547005 | 0.00074041 |
| RAPGEF5 | 1.42756375 | 0.000241616 |
| RARRES1 | 2.42244 | 0.00031792 |
| RASA3 | 1.84563625 | 2.13E-05 |
| RASAL3 | 1.8975995 | 0.0026283 |
| RASD1 | 2.50794225 | 1.10E-05 |
| RASGEF1A | 1.676536 | 0.00141021 |
| RASGRF1 | 3.586074 | 3.26E-05 |
| RASGRP1 | -1.33626125 | 0.000433691 |
| RASGRP4 | 1.7766445 | 0.006032758 |
| RASL10B | -2.34520525 | 0.040401261 |
| RASL11A | 1.34480625 | 0.042606994 |
| RASSF4 | 1.07094725 | 0.005268062 |
| RASSF5 | 1.146216 | 0.005635853 |
| RBBP5 | -1.1033025 | 0.0063118 |
| RBBP8NL | -1.31566725 | 0.048890132 |
| RBFOX3 | -2.7048225 | 0.020809187 |
| RBP4 | 6.59457 | 0.000122748 |
| RBP7 | -2.65574475 | 0.001473632 |
| RCAN1 | 1.10606525 | 0.006033965 |
| RECQL4 | 2.6392785 | 0.034170675 |
| REL | 1.07471825 | 0.005459758 |
| RELB | 1.58555575 | 0.004050039 |
| RETREG1 | 1.14507925 | 0.004365916 |
| RFX4 | 1.46421225 | 0.004495261 |
| RGS1 | 6.107952 | 1.31E-07 |
| RGS10 | 2.02001575 | 0.000518454 |
| RGS11 | 1.5112945 | 0.010225027 |
| RGS13 | 4.71258475 | 1.51E-08 |
| RGS18 | 2.67290125 | 1.06E-05 |
| RGS22 | 2.7276605 | 0.049221321 |
| RGS9BP | -1.499979 | 0.000555385 |
| RHAG | -1.029766 | 0.030932113 |
| RHOB | 2.861296 | 0.000125529 |
| RHOH | 2.63130675 | 0.041458757 |
| RHOU | 1.39765575 | 0.006812251 |
| RIBC2 | 1.21932475 | 0.049460801 |
| RIMS4 | 2.22638025 | 0.004806228 |
| RIN1 | 1.343398 | 0.005625508 |
| RINL | 1.798251 | 0.0026376 |
| RIPK3 | 1.78802875 | 8.10E-05 |
| RIPPLY3 | 9.020111 | 5.65E-11 |
| RIT2 | -3.29668325 | 0.001296734 |
| RMI1 | -1.42015525 | 0.006689911 |
| RNASE10 | 2.2200505 | 8.74E-05 |
| RND1 | 3.26929425 | 0.007497881 |
| RNF149 | 1.07322725 | 0.001567301 |
| RNF224 | -1.98842775 | 0.002209108 |
| RNF39 | 1.466119 | 0.000118297 |
| RNF43 | -1.02449975 | 0.000343539 |
| ROBO1 | 1.63856725 | 1.73E-05 |
| RORA | 1.40459125 | 0.000195098 |
| RPP14 | -1.8620385 | 0.026509853 |
| RPP21 | -1.2331285 | 0.049206308 |
| RRM2 | 1.13669825 | 0.030311765 |
| RSAD1 | -1.16061025 | 0.007478562 |
| RSPH1 | -1.26421475 | 0.013961032 |
| RSPO1 | -2.37229475 | 0.015403475 |
| RTKN2 | 1.306977 | 0.011085454 |
| RTL1 | 1.63245525 | 0.044023821 |
| RTP1 | 1.93872225 | 0.011873962 |
| RTP4 | 1.15235225 | 0.025402094 |
| RUBCNL | 2.27044375 | 0.001802028 |
| RUFY4 | 2.192677 | 0.00227242 |
| RUNX1T1 | -1.241332 | 0.000316074 |
| RUNX2 | 1.463274 | 0.000146422 |
| RWDD2B | -1.45680075 | 0.008637742 |
| RWDD3 | -1.06617775 | 0.000471335 |
| S100A11 | 1.5752235 | 0.000175855 |
| S100A12 | 3.913041 | 0.007375096 |
| S100A2 | 3.495005 | 0.000512057 |
| S100A4 | 2.1218095 | 0.000214284 |
| S100A5 | 2.02043875 | 0.018178686 |
| S100A6 | 1.610224 | 0.015448219 |
| S100A7 | 3.08592975 | 0.004493412 |
| S100A8 | 2.326604 | 0.018359266 |
| S100A9 | 3.01932325 | 0.006051121 |
| SALL4 | -1.84498225 | 0.000543402 |
| SAMD14 | 1.949139 | 0.000925773 |
| SAMD4A | 1.27622925 | 0.001547047 |
| SAMSN1 | 2.42998825 | 2.21E-05 |
| SAP18 | -1.16416 | 0.024942656 |
| SASH3 | 2.59685375 | 0.000502211 |
| SAT1 | 1.44862425 | 0.001138881 |
| SATB2 | -1.31758625 | 0.000391166 |
| SAYSD1 | -1.19255175 | 0.01069697 |
| SBK3 | 2.72365775 | 0.000249541 |
| SBNO2 | 1.36088975 | 0.001254732 |
| SC5D | -1.24109575 | 0.010224691 |
| SCARB1 | 1.403159 | 0.000580963 |
| SCML1 | 1.050959 | 0.000387302 |
| SCML4 | 1.57027975 | 0.023661835 |
| SCN10A | -2.39280775 | 0.001749333 |
| SCN9A | 1.06060725 | 0.007313947 |
| SCNN1G | -1.22063925 | 0.002025881 |
| SDC1 | 1.49068175 | 0.008663054 |
| SDC4 | 1.48557575 | 7.37E-05 |
| SELL | 2.36783425 | 0.000363783 |
| SELPLG | 3.986255 | 3.32E-07 |
| SEMA3A | 2.4848775 | 0.001732998 |
| SEMA4B | 1.0996675 | 0.00093919 |
| SEMA4C | 2.04430775 | 0.001567223 |
| SEMA7A | 1.6116905 | 0.000187875 |
| SEPT3 | 5.153902 | 0.008618242 |
| SERPINA3 | 7.3576715 | 3.20E-05 |
| SERPINB13 | 2.85367125 | 0.012811607 |
| SERPINB2 | 3.444314 | 7.65E-06 |
| SERPINB9 | 1.92997925 | 0.002208098 |
| SERPINE1 | 5.60981225 | 3.06E-05 |
| SERPINF1 | 1.8586 | 0.000118587 |
| SERPING1 | 1.185432 | 0.000573444 |
| SFMBT2 | -1.499608 | 0.004640952 |
| SFRP1 | 1.07812275 | 0.00022842 |
| SFRP2 | 1.12061325 | 0.008609358 |
| SGK1 | 1.60937125 | 0.002945823 |
| SGO2 | -1.584683 | 0.001746783 |
| SH2B2 | 3.1402045 | 0.019509435 |
| SH2D1B | 5.5703515 | 0.000386132 |
| SH2D2A | 2.49432025 | 0.015629274 |
| SH2D6 | 1.45507575 | 0.007426259 |
| SHCBP1 | 2.06395575 | 0.000743139 |
| SHLD1 | -1.5816545 | 0.000974631 |
| SHOC1 | -2.48120325 | 0.00099129 |
| SHOX2 | -3.60452325 | 0.006171695 |
| SHROOM1 | 1.2867835 | 0.032895236 |
| SIGLEC10 | 3.31032775 | 4.51E-05 |
| SIGLEC15 | 5.47979175 | 2.54E-06 |
| SIK1 | 2.795242 | 0.00012052 |
| SIRPB2 | 1.97804725 | 0.016971699 |
| SIT1 | 3.009844 | 0.002024082 |
| SKA1 | 3.52811 | 0.005957927 |
| SKA3 | 2.19095575 | 0.002840171 |
| SKIDA1 | -1.135431 | 0.001291757 |
| SKOR2 | -2.31669675 | 0.025595101 |
| SLA | 3.13032875 | 0.000541027 |
| SLA2 | 3.10976025 | 0.004297667 |
| SLAMF1 | 1.69896475 | 0.00441341 |
| SLAMF6 | 2.185831 | 0.000782542 |
| SLAMF7 | 6.98609725 | 0.000200003 |
| SLAMF8 | 3.3673115 | 5.44E-06 |
| SLC10A1 | -2.3360255 | 0.000332483 |
| SLC10A2 | -1.15112675 | 0.000164935 |
| SLC10A6 | 2.540387 | 0.000509716 |
| SLC11A1 | 3.874287 | 3.50E-06 |
| SLC12A5 | 1.4026605 | 0.031866879 |
| SLC15A3 | 1.05300775 | 0.007471732 |
| SLC16A6 | 2.24744625 | 0.003211509 |
| SLC18A2 | 1.61948475 | 0.000280514 |
| SLC20A1 | 1.6095795 | 2.51E-05 |
| SLC22A20P | -1.95743075 | 0.036874545 |
| SLC24A1 | 1.830482 | 0.030492752 |
| SLC25A25 | 2.646441 | 0.000408312 |
| SLC25A33 | 2.80513025 | 3.23E-07 |
| SLC25A53 | 2.64056075 | 0.005806392 |
| SLC26A1 | -1.5175925 | 1.25E-05 |
| SLC26A3 | -2.72672275 | 0.000776667 |
| SLC26A4 | -1.868275 | 0.000198918 |
| SLC26A5 | 3.31567075 | 0.000774616 |
| SLC27A6 | 5.873768 | 0.003288138 |
| SLC29A2 | 1.9556965 | 0.00070461 |
| SLC2A11 | -1.4387575 | 0.000198671 |
| SLC2A3 | 2.2595415 | 2.66E-05 |
| SLC30A2 | 1.019106 | 0.039101918 |
| SLC35D1 | -1.38246575 | 5.52E-05 |
| SLC36A2 | -2.4220865 | 4.59E-05 |
| SLC37A2 | 6.506521 | 1.82E-08 |
| SLC38A1 | 1.517919 | 0.000217373 |
| SLC38A2 | 1.4567555 | 0.000140388 |
| SLC38A8 | 2.4878215 | 0.003643239 |
| SLC46A2 | 1.5152865 | 0.000526404 |
| SLC4A1 | -1.7011305 | 3.12E-05 |
| SLC4A7 | 1.141947 | 0.032041311 |
| SLC5A5 | 2.269204 | 0.012809891 |
| SLC6A11 | -1.174757 | 0.001724254 |
| SLC6A17 | 1.32786225 | 0.007452055 |
| SLC6A2 | 3.6066625 | 0.03225883 |
| SLC7A1 | 1.43219275 | 0.000721948 |
| SLC7A11 | 1.6530115 | 0.000159033 |
| SLC7A13 | 4.44139625 | 0.032004592 |
| SLC7A5 | 2.6022295 | 2.98E-05 |
| SLC9C1 | -2.37384925 | 4.92E-05 |
| SLCO1A2 | -1.97672275 | 0.013077092 |
| SLCO1B3 | 2.7210425 | 0.013877075 |
| SLCO1C1 | 1.4018325 | 0.016278641 |
| SLCO2A1 | 1.158457 | 0.000201176 |
| SLFN14 | -1.2571535 | 0.006317774 |
| SLITRK6 | -2.9032105 | 0.006513351 |
| SLN | 3.97037425 | 0.004818763 |
| SMC1B | 3.98782125 | 4.75E-05 |
| SMCO3 | 1.75223575 | 0.013596145 |
| SMG5 | -1.0836845 | 0.000668104 |
| SMIM18 | -1.74369475 | 0.00550588 |
| SMIM35 | 2.70743075 | 0.01173786 |
| SMIM5 | -2.13390825 | 4.01E-05 |
| SMLR1 | -1.81377125 | 4.86E-05 |
| SMOX | 1.292019 | 0.005471809 |
| SNAI1 | 1.733122 | 0.000589836 |
| SNAI3 | -1.7924815 | 0.013538637 |
| SNAP25 | 1.9321465 | 0.000551022 |
| SNPH | 1.01581375 | 0.033080798 |
| SNX16 | -1.17341475 | 6.52E-05 |
| SOAT1 | 2.97795425 | 0.000444278 |
| SOCS3 | 4.04110325 | 0.000919184 |
| SORCS2 | 2.2685025 | 0.047142769 |
| SORCS3 | 2.8458765 | 0.00010008 |
| SOX17 | 1.8796415 | 0.002754792 |
| SOX9 | 1.40436825 | 0.009161342 |
| SP5 | 3.58104625 | 0.015278014 |
| SPACA7 | 3.13648225 | 0.011057161 |
| SPAG5 | 1.65132225 | 0.00124932 |
| SPAG6 | 1.5013625 | 0.004906509 |
| SPATA4 | 1.2275395 | 0.02809922 |
| SPATC1 | -1.381183 | 0.046461416 |
| SPC24 | 3.698062 | 0.000172681 |
| SPC25 | 3.5876575 | 0.002385506 |
| SPEF2 | 1.96897575 | 0.042184193 |
| SPHK1 | 1.70837025 | 9.14E-06 |
| SPI1 | 3.494417 | 5.26E-05 |
| SPIC | 2.63306175 | 0.026289298 |
| SPIDR | 1.07057925 | 0.003449091 |
| SPN | 1.8061955 | 0.000145755 |
| SPP1 | 1.01516575 | 0.008385928 |
| SPRED3 | 1.15511175 | 0.010633852 |
| SPRY2 | 1.5081525 | 0.004286288 |
| SPRY4 | 1.42585 | 0.00197769 |
| SPSB1 | 3.53391625 | 9.75E-06 |
| SPTBN5 | 1.6782165 | 0.044792445 |
| SPTLC1 | 1.11489375 | 0.006724113 |
| SPTSSB | -1.5162695 | 0.042883988 |
| SRGN | 1.74043625 | 7.76E-06 |
| SRXN1 | 4.71450375 | 3.66E-05 |
| SSTR5 | -1.72672275 | 0.034764066 |
| ST6GAL2 | 2.81007875 | 0.008688647 |
| ST8SIA1 | -1.19563025 | 0.001727913 |
| STAC3 | 1.43392 | 8.30E-05 |
| STAG3 | 1.169317 | 0.000633648 |
| STAT4 | 5.2883115 | 0.000729983 |
| STC2 | 2.367159 | 0.026896398 |
| STEAP1 | -1.46136325 | 0.000555711 |
| STEAP2 | -1.01171 | 0.021446806 |
| STEAP3 | 1.4010955 | 0.00477606 |
| STEAP4 | 3.97506575 | 0.000664644 |
| STK17B | 1.38160375 | 0.000320392 |
| STK32B | 1.01993175 | 0.022970463 |
| STOM | 1.47827275 | 1.83E-05 |
| STPG2 | -1.5031715 | 0.045136447 |
| STXBP1 | 1.02535575 | 0.000918669 |
| STXBP5L | -2.392709 | 0.034291972 |
| SUSD5 | 2.1573395 | 0.013979334 |
| SV2C | 1.8035535 | 0.030181689 |
| SYCE1L | -1.5336275 | 0.00631144 |
| SYCP2 | 1.4287645 | 0.02128739 |
| SYDE1 | -1.485624 | 0.001117911 |
| SYK | 1.795208 | 0.000329935 |
| SYPL2 | -2.4181065 | 0.004515232 |
| SYT11 | 1.08320275 | 0.003842652 |
| SYT5 | 2.746816 | 0.044647578 |
| SYT6 | 3.519204 | 0.000909352 |
| SYTL3 | 1.97228325 | 0.002128947 |
| SYTL5 | 1.844917 | 0.000364532 |
| TAC4 | 1.84272375 | 0.006051859 |
| TACC3 | 2.3421315 | 1.29E-05 |
| TAF5 | -1.35331325 | 0.006134763 |
| TAFA1 | -3.2991585 | 0.004821802 |
| TAFA3 | 4.2570845 | 5.30E-07 |
| TAFA5 | -1.0770565 | 0.005648745 |
| TAGAP | 2.6594025 | 0.000178197 |
| TAGLN | 1.512582 | 0.002615268 |
| TAL2 | -4.54458875 | 0.011671381 |
| TANGO2 | 1.0265155 | 0.000882837 |
| TAP1 | 1.47808 | 1.50E-05 |
| TBATA | 3.69615875 | 0.000234549 |
| TBC1D10A | 1.03176175 | 0.000462424 |
| TBC1D10C | 1.75118425 | 0.003980131 |
| TBC1D21 | -2.272441 | 0.000776625 |
| TCIRG1 | 1.1554065 | 0.000294354 |
| TCN1 | 5.06516725 | 0.000237262 |
| TCP10 | -1.8729635 | 0.01221658 |
| TCP11 | 2.9104735 | 0.018258702 |
| TCTA | -1.0137045 | 0.000347868 |
| TCTE1 | 2.019204 | 0.013732269 |
| TCTE3 | 1.1455115 | 0.01734058 |
| TDGF1 | 2.7034455 | 3.26E-06 |
| TEKT1 | 3.44615875 | 6.23E-06 |
| TENM1 | -3.4186445 | 9.08E-05 |
| TENT5B | 2.33323675 | 0.000414225 |
| TESMIN | -3.25858525 | 0.001764768 |
| TESPA1 | 4.1982545 | 0.005287204 |
| TET1 | -1.1420685 | 0.047890522 |
| TEX12 | 2.0953315 | 0.000519306 |
| TEX29 | 2.6785615 | 1.35E-05 |
| TEX36 | 1.9767225 | 0.019776033 |
| TEX48 | 1.03531375 | 0.047416865 |
| TF | 2.894081 | 4.79E-05 |
| TFAP2B | -1.24339275 | 0.000103262 |
| TFAP2C | 3.11491025 | 0.047821783 |
| TFCP2L1 | 1.469955 | 0.001609647 |
| TFF3 | 1.46920425 | 0.017384036 |
| TFPI2 | 1.70761725 | 0.000878189 |
| TGFB1 | 1.16394575 | 0.000412841 |
| TGIF1 | 1.0330795 | 0.046851894 |
| THAP9 | -1.230137 | 0.008129858 |
| THBD | 1.934576 | 0.000918865 |
| THBS4 | 2.75595675 | 0.004469823 |
| THEM6 | -1.25738125 | 0.001845401 |
| THSD1 | -1.463756 | 0.000807423 |
| THSD7B | -1.29120575 | 0.001188752 |
| TICAM2 | 2.16602175 | 0.008596488 |
| TIFAB | 2.59824425 | 0.000635298 |
| TIGIT | 2.61278525 | 5.80E-05 |
| TIMD4 | 2.5316925 | 8.34E-05 |
| TIMELESS | 1.2246265 | 0.004052609 |
| TIMP1 | 2.39501575 | 1.67E-05 |
| TIPARP | 1.948389 | 5.13E-06 |
| TK1 | 1.1374505 | 0.008392046 |
| TLR2 | 1.4040565 | 0.003841083 |
| TLR4 | 1.49027225 | 1.84E-05 |
| TLR7 | 1.284725 | 0.00165207 |
| TLR8 | 3.10524075 | 1.25E-06 |
| TMC5 | 1.38466175 | 0.030773502 |
| TMC7 | 1.1838155 | 0.037454735 |
| TMEM132A | 1.22552275 | 9.21E-05 |
| TMEM154 | 1.8620225 | 0.001443955 |
| TMEM156 | 2.93232075 | 0.000615949 |
| TMEM171 | 1.07328325 | 0.00044028 |
| TMEM173 | 1.26174325 | 0.012693027 |
| TMEM178B | 2.05357975 | 0.011347105 |
| TMEM182 | -1.5831685 | 0.002748397 |
| TMEM200A | 1.21172575 | 0.040325243 |
| TMEM221 | 1.673372 | 0.046224286 |
| TMEM229A | 1.339526 | 0.000167386 |
| TMEM239 | 3.2378215 | 0.000166755 |
| TMEM252 | 2.159935 | 0.006473317 |
| TMEM265 | -1.278597 | 0.029397136 |
| TMEM266 | -1.2151605 | 0.024646077 |
| TMEM273 | 2.07665375 | 0.008396322 |
| TMEM40 | 3.291788 | 0.011398025 |
| TMEM45A | 1.828456 | 0.005617364 |
| TMEM59L | 1.79849475 | 0.005905148 |
| TMEM71 | 1.23236975 | 0.000621255 |
| TMEM86A | 1.2811305 | 0.000487193 |
| TMEM89 | -1.0197265 | 0.027254148 |
| TMEM92 | 3.0842915 | 0.010772864 |
| TMPRSS13 | 3.12027625 | 0.00907531 |
| TMPRSS3 | 2.15183275 | 0.016738772 |
| TMPRSS4 | -1.06938825 | 0.02562534 |
| TNF | 4.24269175 | 0.003844557 |
| TNFAIP3 | 3.0663555 | 2.86E-05 |
| TNFAIP6 | 3.82165275 | 0.002824274 |
| TNFRSF12A | 2.98695425 | 9.20E-05 |
| TNFRSF18 | 6.04300875 | 1.03E-08 |
| TNFRSF19 | -1.29868075 | 0.001025422 |
| TNFRSF1B | 3.60453875 | 2.08E-07 |
| TNFRSF4 | 2.488668 | 4.85E-05 |
| TNFSF11 | 3.718724 | 8.32E-06 |
| TNFSF13B | 2.80167325 | 0.021236861 |
| TNFSF14 | 4.19767675 | 0.011420491 |
| TNFSF8 | 2.10496475 | 0.003920989 |
| TNIP3 | 4.532312 | 0.001113015 |
| TNNI1 | 2.1785615 | 0.043873973 |
| TNNT1 | 1.05751725 | 0.035326953 |
| TNR | -2.97916475 | 0.024628703 |
| TOMM20L | 3.771017 | 6.70E-05 |
| TOP2A | 3.35060275 | 0.000528388 |
| TOX2 | 3.29103875 | 1.51E-05 |
| TP53INP1 | 3.98794575 | 2.01E-06 |
| TP53TG5 | 1.056304 | 0.012525116 |
| TP73 | -1.39414 | 0.039130969 |
| TPH1 | 2.37296325 | 0.011722234 |
| TPM4 | 1.15267875 | 0.001651395 |
| TPO | 1.66864475 | 0.002937301 |
| TRAF1 | 1.229454 | 0.0010479 |
| TRARG1 | -2.188722 | 0.024108905 |
| TREM1 | 3.25265125 | 0.001850845 |
| TREM2 | 1.82185475 | 0.013637321 |
| TRIL | -1.01474925 | 0.013344317 |
| TRIM13 | -1.048585 | 0.00059979 |
| TRIM31 | 2.21065525 | 0.000257949 |
| TRIM36 | 1.25292925 | 0.037061702 |
| TRIM63 | 1.0249175 | 0.016861411 |
| TRIM72 | 4.749912 | 1.75E-08 |
| TRMT5 | -1.040347 | 0.000294378 |
| TRNAK-UUU_8 | -1.14624075 | 0.009440825 |
| TRNAN-GUU | 1.47672275 | 0.001433234 |
| TRNAN-GUU_12 | 1.2886445 | 0.014924823 |
| TROAP | 4.7276375 | 0.000481503 |
| TRPC3 | -1.21247825 | 0.021551985 |
| TRPC6 | 1.253958 | 0.007188026 |
| TRPM2 | 5.494723 | 1.66E-05 |
| TRPM3 | -1.425429 | 8.04E-05 |
| TRPM5 | 4.67654325 | 3.86E-06 |
| TRPV2 | 3.99743275 | 3.83E-05 |
| TSC22D3 | 1.6680635 | 0.004451645 |
| TSEN54 | -1.2850765 | 0.004403532 |
| TSHR | -1.63701825 | 0.003940925 |
| TSKU | 3.04995875 | 1.08E-06 |
| TSPAN32 | 2.053676 | 0.010233916 |
| TSPYL1 | 1.19091525 | 0.038405802 |
| TSSK4 | 3.4841485 | 9.48E-05 |
| TTBK1 | 1.15623375 | 0.016492238 |
| TTC21A | 1.05604425 | 0.038780425 |
| TTC25 | 2.0084075 | 0.003031079 |
| TTC39A | 2.112354 | 0.000649399 |
| TTK | 2.41933175 | 0.00259936 |
| TTLL7 | 1.74921825 | 7.38E-05 |
| TTYH1 | -1.1018755 | 0.004379313 |
| TUBAL3 | -3.43517625 | 0.000114192 |
| TUBB1 | -1.44624475 | 0.009445756 |
| TUBB4A | 1.22052875 | 0.001927484 |
| TXK | 1.3216585 | 0.026447486 |
| TXNDC2 | 1.06575875 | 0.042436126 |
| TXNIP | 1.32353 | 7.44E-05 |
| TYR | -3.14079525 | 0.035917618 |
| TYROBP | 3.00630075 | 6.12E-05 |
| UBAP1L | 1.80170925 | 0.024337269 |
| UBASH3A | 2.26986525 | 0.00050467 |
| UBASH3B | 2.6015775 | 2.79E-06 |
| UBE2C | 3.15024475 | 0.001343973 |
| UBXN10 | 3.93689075 | 0.000700712 |
| UCHL1 | 2.7715495 | 0.009445046 |
| UGT8 | -1.35968575 | 0.002034888 |
| UHRF1 | 2.440573 | 1.51E-05 |
| UNC13C | -1.422333 | 0.004960742 |
| UNC93B1 | 2.23518 | 0.000309869 |
| UNCX | -2.43386675 | 0.00103616 |
| USP2 | 2.17021775 | 1.50E-05 |
| USP36 | 1.25221 | 0.000643717 |
| VASH2 | -1.4914955 | 0.000124416 |
| VAV1 | 2.6719495 | 6.15E-05 |
| VCAM1 | 1.029771 | 0.000984832 |
| VGF | 2.22581975 | 0.022495047 |
| VIM | 1.89240925 | 0.000323653 |
| VKORC1 | -1.3898065 | 0.001080209 |
| VMO1 | -1.01268775 | 0.00041208 |
| VOPP1 | 1.05665025 | 0.000701791 |
| VSIG4 | 2.11086175 | 0.001460334 |
| VSIR | 1.2404315 | 0.000453039 |
| VSNL1 | -1.07262875 | 0.00122968 |
| VSTM1 | 4.359034 | 0.002014503 |
| VSX1 | 1.393222 | 0.014343938 |
| VTCN1 | 1.87372575 | 0.005882975 |
| VWA3A | 2.233857 | 0.000459046 |
| WAS | 1.50569075 | 0.002234123 |
| WBP1L | 1.2862015 | 0.000810659 |
| WDCP | -1.192861 | 5.73E-05 |
| WDFY4 | 2.9416785 | 0.000955542 |
| WDHD1 | 1.05845625 | 0.003794135 |
| WDR4 | 1.0583315 | 0.003897775 |
| WDR62 | 1.04637675 | 0.029600925 |
| WIF1 | 1.67969775 | 0.023676747 |
| WIPF1 | 1.1798785 | 0.006415831 |
| WNK4 | -1.282671 | 3.95E-05 |
| WNT10A | 3.38636175 | 0.008259605 |
| WNT10B | 5.656884 | 2.10E-05 |
| WNT2 | 4.7778465 | 0.001303548 |
| WNT8B | 3.74100775 | 1.77E-07 |
| WSCD1 | -1.92548925 | 0.000101966 |
| XCR1 | 4.90998525 | 0.001740621 |
| XYLB | -1.115497 | 0.005978252 |
| YPEL4 | 2.41708025 | 0.003860973 |
| ZBED1 | -1.2877185 | 0.003607889 |
| ZBED8 | -1.2324615 | 0.00344077 |
| ZBTB12 | -1.58052225 | 0.000108235 |
| ZBTB16 | 4.770706 | 8.32E-06 |
| ZBTB32 | 2.0499765 | 0.041239187 |
| ZBTB45 | -1.029489 | 0.002378189 |
| ZC3H12A | 1.7343775 | 0.001827604 |
| ZC3H8 | -1.072083 | 0.006830324 |
| ZCCHC12 | 2.03232075 | 0.044338861 |
| ZCWPW1 | 2.89056075 | 0.000206969 |
| ZFAND5 | 2.99329325 | 2.57E-07 |
| ZFP30 | -1.1913305 | 0.001366058 |
| ZFP36 | 2.10148925 | 0.003633783 |
| ZFP36L2 | 1.219379 | 3.44E-05 |
| ZFP57 | 1.86470125 | 0.002398472 |
| ZFP69B | -1.32350775 | 0.003121915 |
| ZFPM1 | -1.14086475 | 0.000900452 |
| ZIM2 | -1.067393 | 0.049925423 |
| ZMYM1 | -1.32713075 | 0.000214991 |
| ZMYND19 | -1.01291425 | 0.001919378 |
| ZNF135 | -1.39677775 | 0.01079306 |
| ZNF16 | -1.677911 | 0.027333832 |
| ZNF217 | 1.25819825 | 6.48E-05 |
| ZNF275 | 1.26116375 | 0.001474846 |
| ZNF341 | -1.175698 | 0.000647725 |
| ZNF385A | 1.58559125 | 0.002959897 |
| ZNF397 | -1.00440275 | 0.001210304 |
| ZNF398 | -1.173682 | 0.014319743 |
| ZNF420 | -1.1983805 | 0.000416162 |
| ZNF449 | -1.136886 | 0.036079083 |
| ZNF510 | -1.15513375 | 0.00176239 |
| ZNF514 | 1.03888375 | 0.000362866 |
| ZNF526 | -1.51487425 | 0.004048529 |
| ZNF575 | 2.6229635 | 0.01618672 |
| ZNF628 | 1.2738785 | 0.003754781 |
| ZNF658 | -1.38553625 | 0.013669609 |
| ZNF668 | -1.45026775 | 0.011384125 |
| ZNF697 | -1.2956385 | 1.38E-05 |
| ZNF774 | -1.03673325 | 0.019198424 |
| ZNF782 | -1.06527525 | 0.001241277 |
| ZNF789 | -1.3037365 | 0.016083651 |
| ZNF831 | 3.16506275 | 0.022567259 |
| ZNF862 | -1.398297 | 0.027277598 |
| ZPBP2 | 2.08496275 | 0.012819002 |
| ZSCAN2 | -1.4484045 | 0.001565078 |
| ZSWIM1 | -1.084663 | 9.08E-05 |
| ZSWIM4 | 2.042805 | 0.002710733 |
| ZSWIM5 | -1.01806975 | 6.60E-05 |
| ZWILCH | 2.089105 | 0.000139611 |
